# circFL-seq reveals full-length circular RNAs with rolling circular reverse transcription and nanopore sequencing

**DOI:** 10.1101/2021.07.05.451107

**Authors:** Zelin Liu, Changyu Tao, Shiwei Li, Minghao Du, Yongtai Bai, Xueyan Hu, Yu Li, Jian Chen, Ence Yang

## Abstract

Circular RNAs (circRNAs) act through multiple mechanisms with their sequence features to fine-tune gene expression networks. Due to overlapping sequences with linear cognates, identifying internal sequences of circRNAs remains a great challenge, which hinders comprehensive understanding of circRNA functions and mechanisms. Here, based on rolling circular reverse transcription (RCRT) and nanopore sequencing, we developed circFL-seq, a full-length circRNA sequencing method, to profile circRNA at the isoform level. With a customized computational pipeline *circfull* to directly identify full-length sequences from rolling circular reads, we reconstructed 77,606 high-quality circRNAs from seven human cell lines and two human tissues. Benefiting from rolling circles and long-read sequencing, circFL-seq showed more than tenfold enrichment of circRNA reads and advantages for both detection and quantification at the isoform level compared to short-read RNA sequencing. The concordance of RT-qPCR and circFL-seq results for the identification of differential alternative splicing suggested wide application prospects for functional studies of internal variants in circRNAs. Moreover, the detection of cancer-related fusion circRNAs at the omics scale may further expand the application of circFL-seq. Together, the accurate identification and quantification of full-length circRNAs make circFL-seq a potential tool for large-scale screening of functional circRNAs.

## Introduction

CircRNAs, a class of covalently closed RNA molecules formed via back-splicing (BS) or lariat precursors, are involved in various biological processes and pathogenesis by fine-tuning the eukaryotic gene regulatory network^1,2^. CircRNAs directly or indirectly regulate target gene expression via diverse mechanisms, which are largely determined by circRNA sequence features. For example, circRNAs may complementarily bind to miRNAs as sponges^3,4^; circRNAs may interact with proteins as scaffolds^5^ or structural components^6^ based on sequence motifs, and several circRNAs are also able to translate into peptides^7,8^ through internal ribosome entry sites (IRESs). Thus, full-length sequences of circRNAs have become the foundation for ascertaining their biological functions in transcriptional plasticity and complexity.

By detecting the back-splicing junctions (BSJs) of circRNAs with deep sequencing, short-read RNA sequencing discriminates circRNAs with low expression (as low as 1% polyadenylated RNA^9^) from their linear cognates. With the benefits of bioinformatics approaches, the full-length sequence of short circRNAs (< 500 nt) could be inferred from a patchwork of BSJs and short fragments^10,11^. However, the full understanding of circRNA isoforms is impossible by using short reads. Single-molecule long-read sequencing has shown methodological advances in identifying circRNAs at the isoform level. Pacific Biosciences (PacBio) sequencing was applied to PCR products for target full-length circRNA sequences in a low-throughput and high-cost way^12^. In contrast, rolling circular amplification (RCA) for circular cDNA of circRNA followed by Oxford Nanopore Technologies (ONT) sequencing can be used to reconstruct genome-wide full-length circRNA^13^. However, the method is limited by the use of ligation, which may also generate circular cDNA from linear residuals. Moreover, the low-coverage reads of full-length circRNAs are more likely to generate false discoveries, quantification bias, and high costs. Thus, an accurate but affordable method to detect full-length circRNA remains to be developed for wide application in screening functional circRNAs at the omics scale.

Because of their native circular structure, reverse transcription of circRNA can produce rolling circles of full-length circRNA sequences, i.e., rolling circular reverse transcription (RCRT), which has been leveraged to distinguish internal sequences of target circRNAs from linear and circular cognates^14,15^. Here, we developed a high-throughput circRNA sequencing method, termed circFL-seq, with RCRT and nanopore long-read sequencing for the identification of circRNA at the isoform level. With a customized computational pipeline *circfull*, we identified 77,606 high-quality full-length circRNA isoforms from seven cell lines and two human tissues. We validated circFL-seq for full-length circRNA detection and quantification by comparison to annotated circRNAs, RNA-seq, isoCirc, and RT-qPCR results. By providing full-length circRNA sequences, circFL-seq allows the study of sequence features, alternative splicing, and differential expression at the isoform level. In addition, circFL-seq also showed the ability to detect fusion circRNAs (f-circ), which were further experimentally validated. Taken together, benefiting from RCRT and long-read sequencing, circFL-seq with higher read coverage has shown advantages for the identification of high-quality full-length circRNAs at a low cost.

## Results

### Sequencing full-length circRNA with circFL-seq

We developed a circFL-seq approach that utilized RCRT with long-read sequencing for full-length circRNA profiling (Figure 1a). First, circRNA was enriched by rRNA depletion, poly(A) tailing (to increase the efficiency of RNase R^16^), and RNase R treatment. Then, first-strand cDNA was synthesized with random primers (P1-N_6_) by RCRT for circRNA and regular reverse transcription for linear residuals. After tailing poly(A) of the first strand and synthesizing the second strand with an anchor primer (P2-T_24_), the double-strand cDNA was amplified with P1 and P2 to construct the final circFL-seq library. To benchmark full-length circRNA sequences of the circFL-seq library, eight known circRNAs were selected for PCR validation in the HEK293T library. Rolling circles of target circRNA (Figure 1b) and the full-length sequences obtained with Sanger sequencing (Figure 1–figure supplement 1) indicated that the circRNAs were successfully amplified. Finally, the circFL-seq library was sequenced by long-read sequencing on the ONT platform.

**Figure 1.**
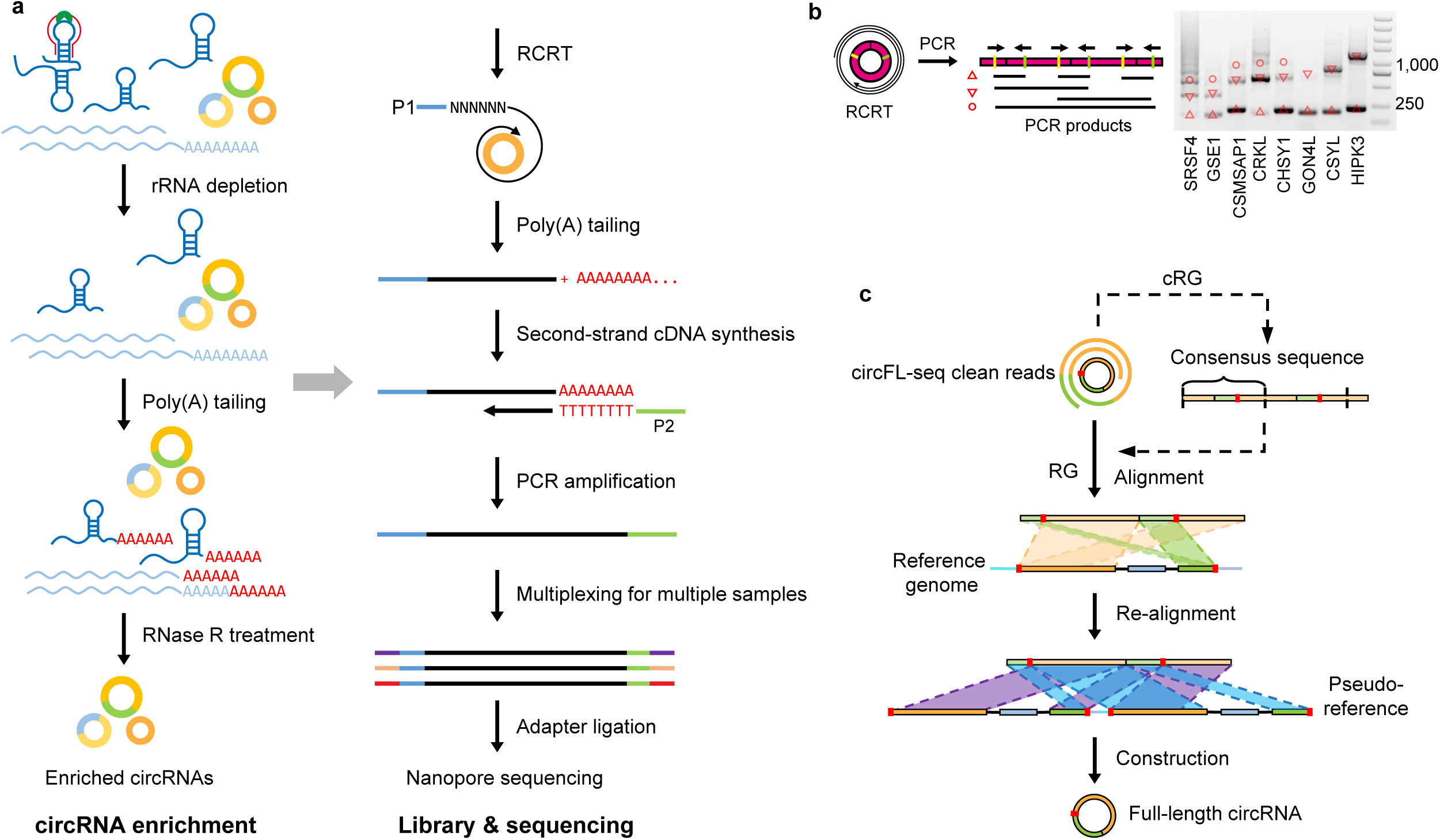
Diagram of circFL-seq workflow. **a.** Experimental operation of circFL-seq consisted of circRNA enrichment, library construction, and nanopore sequencing. **b.** PCR validation of rolling circle products from the circFL-seq cDNA library. The yellow and green lines indicate the positions of the PCR primers. The upward triangle, downward triangle, and circle symbols denote the 0-circle, 1-circle, and 2-circle cDNA products. **c.** Computational pipeline of *circfull*. circFL-seq clean reads were directly used in RG mode or were self-corrected for consensus sequences in cRG mode to reconstruct full-length circRNAs.

Based on the nature of RCRT products in circFL-seq libraries, full-length sequences of circRNAs were reconstructed and structurally annotated with a customized pipeline *circfull*. For the reference guide (RG) mode of *circfull*, clean reads (qscore ≥ 7) were aligned to the reference genome to identify potential back-spliced junctions (BSJs) by the sign of chiastic overlapping segments (Fig 1c). Then, reads were realigned to pseudo-references, generated by concatenating two sequences of potential BSJ regions, to accurately localize circRNA BSJs. In addition, full-length structures were adjusted with multiple alignments from rolling circular segments to improve the construction. To complement the RG mode for low-quality reads, the cRG mode of *circfull* identified consensus sequences (CS) from clean reads with two or more rolling circles, and triply duplicated CS were used as query sequences in RG mode to improve circRNA detection.

### circRNA profiling of eight libraries

To assess the performance of circFL-seq, we applied circFL-seq to eight libraries of six human cell lines (two replicates from HeLa cells, two replicates from SKOV3 cells, one from MCF7 cells, one from VCaP cells, one from SH-SY5Y cells, and one from HEK293T cells) and produced 30 M clean reads sequenced by using one PromethION cell line (Figure 2–figure supplement 1a). We detected 197,252 isoforms of 162,409 circRNA BSJs from 1.3 M reads that contained full-length circRNA sequences (Figure 2a; Figure 2–figure supplement 1b; Figure 2–figure supplement 2a; Supplementary file 1), 2% of which were additionally detected by cRG mode, which refined low-quality reads with CS (Figure 2–figure supplement 2b). Quantification of replicate samples at both the BSJ and isoform levels was highly consistent (Pearson’s *r* > 0.93; Figure 2a,b; Figure 2–figure supplement 3).

**Figure 2.**
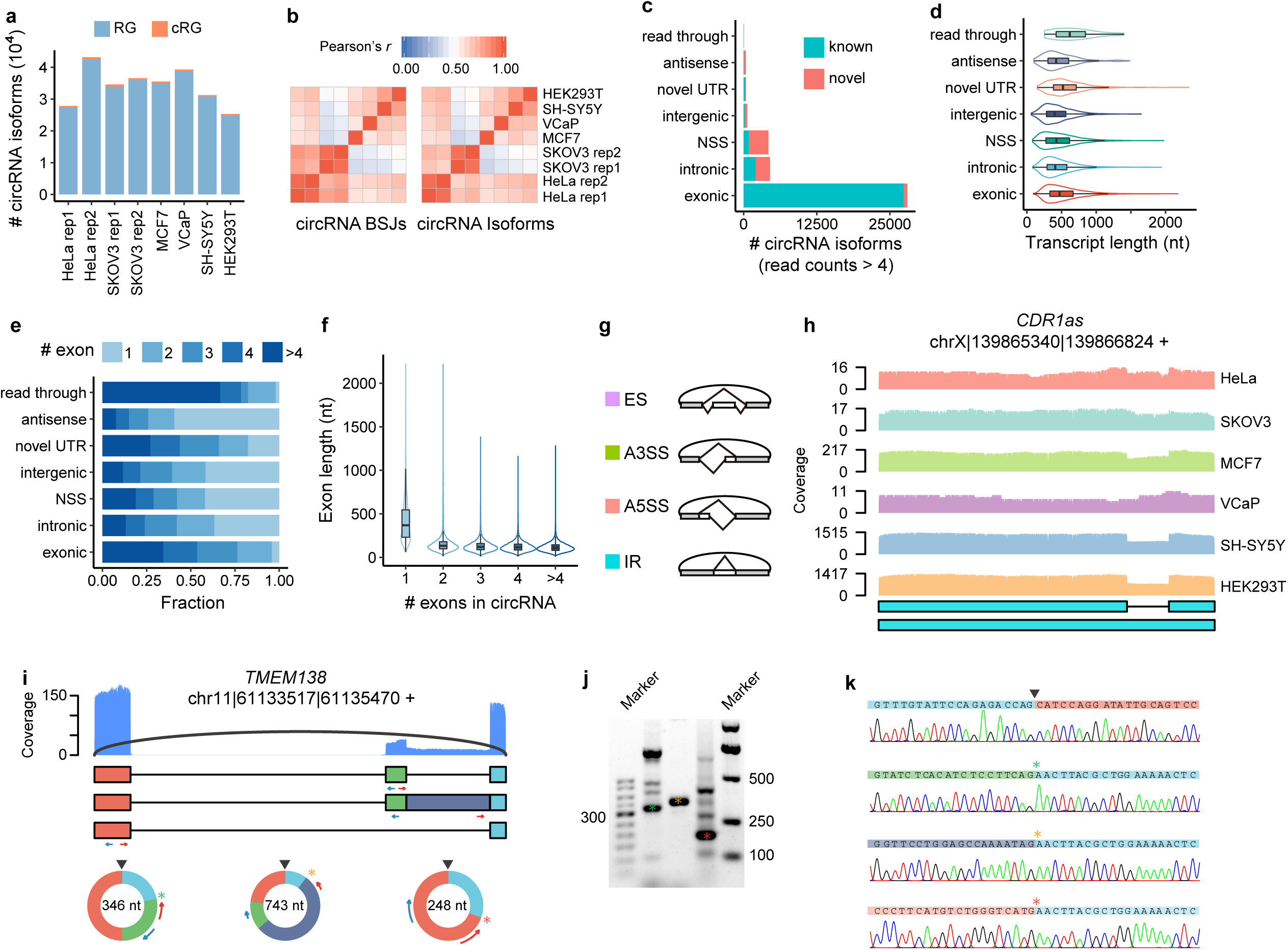
Analysis of full-length circRNA in eight samples. **a.** The stacked bar plot represents the number of full-length circRNA isoforms detected by RG and cRG for six cell lines. **b.** Expression correlation matrix for circRNA BSJs and isoforms of all samples. The color scale corresponds to Pearson’s correlation coefficient. **c.** Stacked bar plot represents the number of circRNA isoforms with read counts ≥5 from known or novel BSJs based on the circRNA database. **d.** Boxplot showing the length distribution per isoform for circRNA isoforms with read counts ≥5 in all samples. Box lefts or rights are lower or upper quartiles, the bar is the median and whiskers are the median ± 1.5× interquartile range. **e.** Stacked bar plot showing the fraction of exon numbers per isoform for circRNA isoforms with read counts ≥5 in all samples. **f.** Boxplot showing the length distribution per exon for circRNA isoforms with read counts ≥5 in all samples. Box bottoms or tops are lower or upper quartiles, the bar is the median, and the whiskers are the median ± 1.5× interquartile range. **g.** Diagram of four types of alternative splicing (AS) events in circRNA: exon skipping (ES), alternative 3’ splice site (A3SS), alternative 5’ splice site (A5SS), and intron retention (IR). **h.** Plot showing the coverage of full-length circRNA reads in the position of *CDR1as* for circFL-seq data of six cell lines (replicate data were merged). Structures of the two isoforms of *CDR1as* are shown at the bottom. **i-k.** AS events (one ES and one IR) of circ-*TMEM138* detected by circFL-seq (**i**), agarose gel electrophoresis (**j**), and Sanger sequencing (**k**). Red/blue arcs are forward/reverse primers for validation of back-splicing junctions (BSJs) and forward splicing junctions (FSJs). Asterisks denote FSJs. Downward triangles denote BSJs.

Based on BSJs and boundary exons identified by circFL-seq, we classified circRNAs into seven types: exonic, intronic, novel splicing site (NSS), intergenic, novel UTR, antisense, and read through (Figure 2–figure supplement 4). For high-quality circRNA isoforms (read counts ≥ 5), the exonic type accounted for 73.2% of the identified circRNA species, while only 0.1% were from read throughs (Figure 2c). Most exnoic types (99.8%) could be verified in at least one database^17–22^ (Figure 2c), which could be attributed to their higher expression levels (Figure 2–figure supplement 5). The median length of each type varied from 402 nt of the intergenic type to 618 nt of read-throughs (Figure 2d), indicating that a notable proportion of circRNAs > 500 nt cannot be reconstructed by RNA-seq. Read-through circRNAs contained more exons (Figure 2e), while the exon length of single-exon circRNA was significantly longer than that of multiple-exon circRNA (*P* < 2.2 × 10^−16^; Figure 2f).

From 65,656 isoforms of 35,251 high-quality circRNA BSJs (read counts ≥ 5), we identified 23,267 internal AS events (Supplementary file 2), including 44.3% exon skipping (ES), 28.1% alternative 3’ splicing site (A3SS), 24.1% alternative 5’ splicing site (A5SS), and 3.5% intron retention (IR) events (Figure 2g). Specifically, IR events increased circRNA length by 289 nt on average (from 363 to 652 nt), which may influence their cellular localization^23^. circFL-seq accurately detected the full length of two isoforms (1485 and 1301 nt) generated by the IR event of *CDR1as*^24^ (Figure 2h). We further experimentally verified 13 AS events (5 of ES, 3 of A3SS, 1 of A5SS, and 4 of IR) of 23 isoforms from 11 high-quality circRNAs in the HeLa cell line. With divergent primers to amplify both BSJs and alternative spliced forward splicing junctions (FSJs), all but a A3SS event were validated by Sanger sequencing (Figure 2i-k; Figure 2–figure supplement 6-8).

### Comparison with RNA-seq and isoCirc for circRNA detection

Based on read-spanning BSJs, short-read sequencing has long been used for genome-wide characterization of circRNAs. Thus, for comparison, eight RNA-seq libraries (150 bp × 2) with the same circRNA enrichment method for the same six cell lines (Pearson’s *r* > 0.90 for replicates, Figure 3–figure supplement 1) were generated and sequenced by the Illumina HiSeq X Ten platform. A total of 95,371 circRNA BSJs were detected by CIRI2^25^ (Supplementary file 3). BSJ reads accounted for 0.15%–0.32% of all RNA-seq reads, an amount ten times lower than that of full-length circRNA reads with circFL-seq (2.2%–8.5%). Compared to known BSJs that have been annotated in the database, the proportion of overlapping BSJs identified by RNA-seq seems to be larger than that identified by circFL-seq (76.7% vs. 40.3%) (Figure 3–figure supplement 2). However, when focusing on high-quality BSJs (read counts ≥ 5), the proportion obtained with circFL-seq (78.9%) was similar to that obtained with RNA-seq, indicating that more reads are required to confidently identify circRNAs by ONT (Figure 3–figure supplement 3). As expected, the full-length construction by RNA-seq was highly dependent on circRNA length (Figure 3–figure supplement 4). Approximately 96.3% of full-length circRNAs reconstructed by RNA-seq were less than 500 nt, while 44.4% of circRNA isoforms detected by circFL-seq were more than 500 nt in length.

A recent method, isoCirc, identified full-length circRNAs by rolling circle amplification (RCA) followed by nanopore sequencing. With a single circFL-seq library of the HEK293 cell line sequenced by MinIon, we compared the features of the RCRT strategy and RCA strategy. circFL-seq produced more full-length circRNA reads per 10^9^ raw bases than isoCirc (11,791 vs. 5820) (Figure 3–figure supplement 5a-c) but identified fewer circRNA isoforms (Figure 3–figure supplement 5d), which resulted in higher read coverage of detected circRNAs (Figure 3–figure supplement 5e). For both methods, circRNAs with higher read counts were more likely to be included in the database (Figure 3–figure supplement 5f-i). When focusing on the top 100 expressed BSJs identified by each method, BSJs from both circFL-seq and RNA-seq (collected from isoCirc study) were annotated in databases, while 22 of the top 100 BSJs identified by isoCirc lacked support from the database, RNA-seq, or circFL-seq results (Figure 3–figure supplement 5m). Specifically, two products were from histone genes whose linear RNAs are resistant to RNase R degradation^16^. At the isoform level in the HEK293 cell line, 87.0% of the common BSJs (10,702/12,307) between circFL-seq and isoCirc identified at least one isoform. Inconsistent isoforms usually had lower read counts with both circFL-seq and isoCirc (Figure 3–figure supplement 5n,o), suggesting again that accurate construction of full-length circRNA requires higher read coverage.

We then compared circFL-seq to isoCirc in two normal human tissues (brain and testis) with high sequencing depth. The saturation curve showed BSJ detection of circFL-seq near saturation, detecting 64,716 BSJs and 40,936 high-quality BSJs (read counts ≥ 5) (Figure 3–figure supplement 6a,b). In contrast, isoCirc identified 141,459 BSJs and 17,801 high-quality BSJs, which was far from saturation (Figure 3–figure supplement 6c,d). The overlapping BSJs were significantly more highly expressed than isoCirc-only BSJs (Figure 3–figure supplement 6e,f), indicating that isoCirc is able to detect more circRNAs with low expression and requires a vast sequencing depth to identify high-quality BSJs as a trade-off (Figure 3–figure supplement 6g-k).

### Evaluation of quantification of full-length circRNAs

circFL-seq better produced circRNAs with high read coverage, which is beneficial for circRNA quantification. For BSJs identified by circFL-seq and RNA-seq, the amounts between the two methods were significantly correlated (Pearson’s *r* = 0.41–0.68, Figure 3a; Figure 3–figure supplement 7). We then detected differentially expressed circRNAs (DECs) between HeLa and SKOV3 cell lines as a test. For highly expressed BSJs (read counts ≥ 10 in at least two samples and detected by both methods), the fold changes of HeLa to SKOV3 cells were highly concordant with RNA-seq results (Pearson’s *r* = 0.78, Figure 3b). By using DESeq2^26^, circFL-seq detected 89 DECs, 58 of which were also identified by RNA-seq. We next selected 16 circRNAs with a wide range of fold changes (7 downregulated, 5 stable, 4 upregulated in the HeLa cell line) to validate the DECs via RT-qPCR. The consistent circFL-seq and RT-qPCR results for both total RNA (Figure 3–figure supplement 8a-c) and RNase R-treated (Figure 3c,d) samples further supported the capabilities of BSJ quantification.

**Figure 3.**
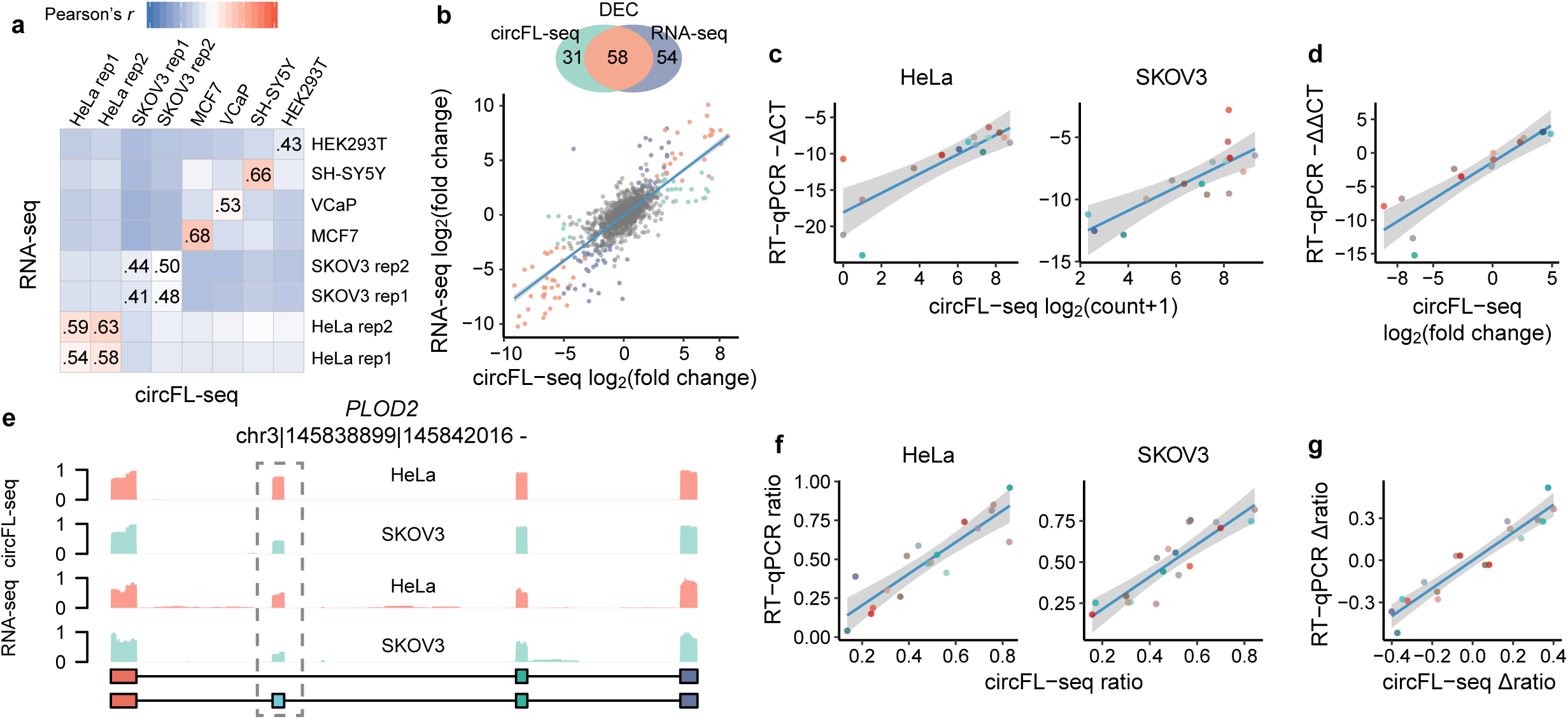
Quantification of circRNA at the BSJ and isoform levels. **a.** Expression correlation matrix of circRNA BSJ quantified by circFL-seq and RNA-seq for six cell lines. The numbers in the matrix represent Pearson’s correlation coefficients. **b.** Comparison of differentially expressed circRNA (DEC) detection between circFL-seq and RNA-seq. Top panel: Venn diagram showing the number of DECs detected by circFL-seq (green), RNA-seq (purple), and both methods (orange). Bottom panel: scatter plot showing the correlation of fold change (log base 2) for HeLa and SKOV3 cells between circFL-seq and RNA-seq. **c.** Scatter plot showing the correlation of the expression levels of 16 circRNA BSJs for HeLa (left) and SKOV3 (right) cells between circFL-seq and RT-qPCR. **d.** Scatter plot showing the correlation of fold changes (log base 2) of the 16 BSJs for HeLa and SKOV3 cells between circFL-seq and RT-qPCR. **e.** Plot showing the coverage of full-length circRNA reads and RNA-seq reads in the position of circRNA from *PLOD2*. The circular structures of the two circRNA isoforms are shown in the lower panel. **f.** Scatter plot showing the correlation of the transcript ratio of 18 circRNA isoforms from 9 circRNA BSJs (each BSJ has two isoforms) for HeLa (left) and SKOV3 (right) cells between circFL-seq and RT-qPCR. The relative expression of target BSJs/isoforms quantified by RT-qPCR was determined with RNase R-treated samples and GAPDH from total RNA without RNase R treatment as a reference. **g.** Scatter plot showing the correlation of the differential ratio (Δratio) of the 18 isoforms for HeLa and SKOV3 cells between circFL-seq and RT-qPCR.

We next evaluated the quantification performance of circFL-seq for circRNA AS. From 87 BSJs that had at least two isoforms detected by both HeLa and SKOV3 cell lines, 193 isoforms were used to quantify the internal variants by the ratio of transcripts (target isoforms to total isoforms from the same BSJ). A total of 90 transcripts had more than 0.1 ratio differences between HeLa and SKOV3 cell lines. For example, circ-*PLOD2* showed a higher ratio of transcripts without ES in the HeLa cell line (0.83 to 0.45) (Figure 3e). Although RNA-seq data also detected differential read coverage at the skipped exon of *PLOD2*, short reads were unable to discriminate different sources of ES events, i.e., linear or circular isoforms. We selected 18 isoforms of 9 circRNA BSJs from a wide range of ratio differences to perform experimental validation. Both the transcript ratios and ratio differences were consistent with the RT-qPCR results (Figure 3f, g; Figure 3–figure supplement 8d-f), supporting the advantage of circFL-seq for quantification at the isoform level.

### circFL-seq reveals fusion circRNAs with long-read sequencing

Fusion circRNA (f-circ) is a specific type of circRNA containing two fusion junctions (one junction may come from gene fusion and the other may be produced by BS) and has been found to play important roles in cancer^27^. Although short reads of RNA-seq may detect both fusion junctions separately, they usually fail to detect f-circ for the short read that is not able to combine two fusions in one fragment. circFL-seq is able to detect f-circ by *circfull* with rolling circular reads separately mapped to two loci in different chromosomes or a > 1 Mbp distance in the same chromosome (Figure 4a). With circFL-seq data, we identified six high-quality f-circ isoforms (read count ≥ 5) affiliated with two fusion genes in the MCF7 cell line. For the five isoforms fused by *GBF1* and *MACROD2* (Figure 4b), we selected two major isoforms (circ_290 nt for 35% and circ_408 nt for 26%) to perform full-length validation. As expected, more than one product was observed (Figure 4c), and the rolling circles of full-length sequences were validated by Sanger sequencing (Figure 4d, Figure 4–figure supplement 1). For comparison, we also used linear RNA from the MCF7 cell line and RNase R-treated RNA from the HeLa cell line as templates. Intriguingly, we validated the fusion junction of E2-E4 with linear RNA as a template (Figure 4c), suggesting the existence of linear RNA production from the fusion gene. The decreased expression of junction E2-E4 after RNase R treatment also supported the linear production, which further suggested that the fusion direction is *GBF1* to *MACROD2* (Figure 4e). In addition, the f-circ fused by two antisense genes (*PRICKLE2-AS1* and *PTPRT-AS1*) was experimentally validated (Figure 4f; Figure 4–figure supplement 2). All the major fusion junctions of *GBF1*:*MACROD2* and E3-E1 of *PRICKLE2-AS1*:*PTPRT-AS1* were validated by RNA-seq of the MCF7 cell line (Figure 4g).

**Figure 4.**
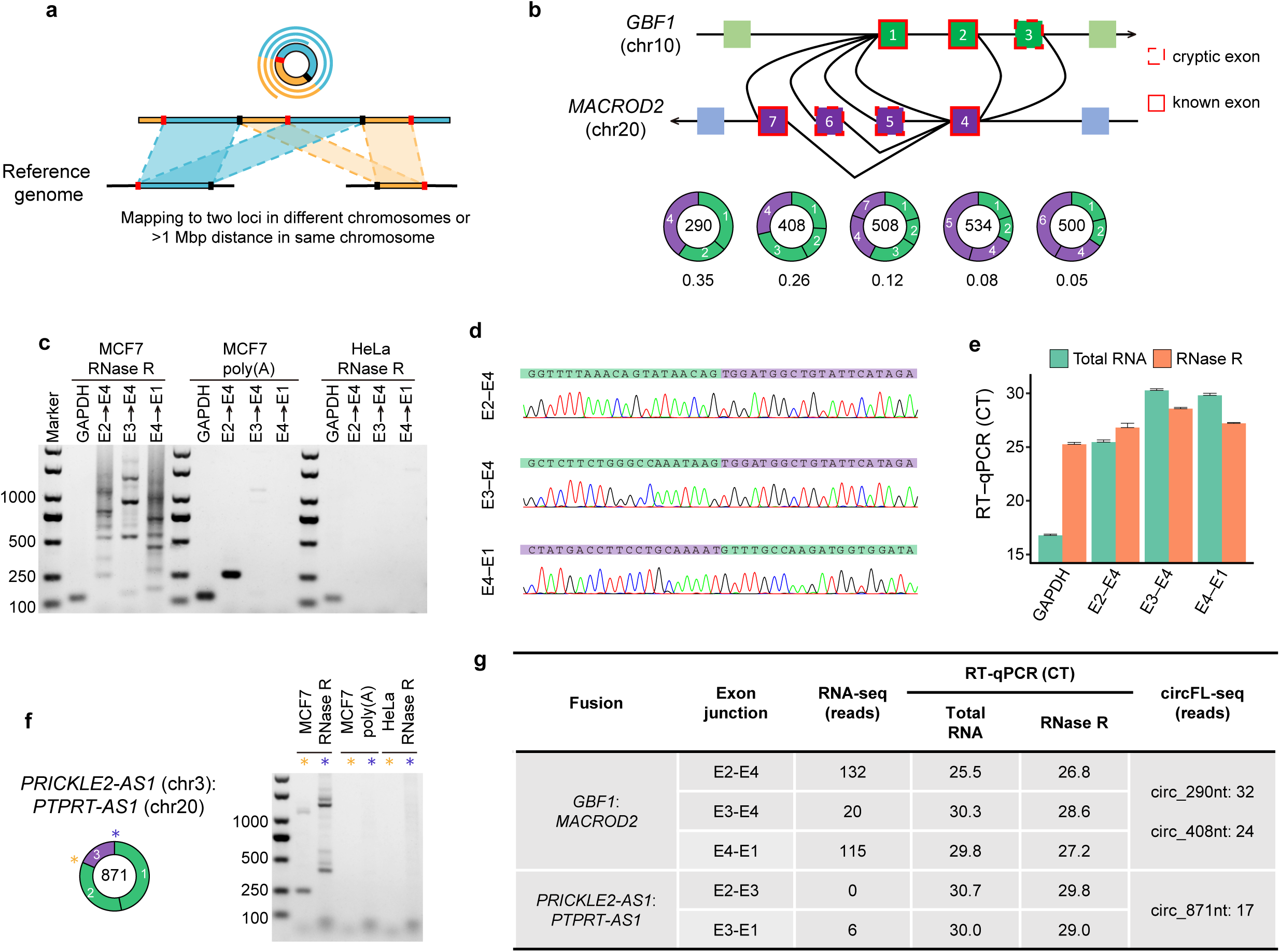
Detection and validation of fusion circRNA (f-circ) in the MCF7 cell line. **a.** Diagram of *circfull* identification of f-circ with circFL-seq data. **b.** Diagram of five high-quality f-circ isoforms (read counts ≥5) fused by *GBF1* and *MACROD2*. The transcript ratio represents the fractions of the isoforms. **c-e** Validation of f-circ junctions from *GBF1*:*MACROD2* by agarose gel electrophoresis (**c**), Sanger sequencing (**d**), and RT-qPCR (**e**). **c.** Agarose gel electrophoresis showing the RT-PCR products of f-circ junctions with RNase R-treated MCF7 and HeLa RNA and poly(A) selected MCF7 RNA as a template. **f.** Agarose gel electrophoresis showing the RT-PCR products of f-circ junctions from *PRICKLE2-AS1:PTPRT-AS1*. **g.** Information on five f-circ junctions detected by circFL-seq, RNA-seq, and RT-qPCR.

## Discussion

In this study, we established a full-length circRNA sequencing method, circFL-seq, by nanopore sequencing. Compared to short-read sequencing, which is limited to reconstructing short circRNAs (< 500 nt), long-read sequencing of circFL-seq has shown advances in comprehensively identifying full-length circRNAs of all sizes (64 nt to 2334 nt in our data and more than 40% identified isoforms > 500 nt). Very recently, isoCirc^13^ and CIRI-long^28^ also employed nanopore sequencing to identify full-length circRNAs by utilizing rolling circles. Both circFL-seq and CIRI-long employed RCRT to produce rolling circles during first-strand synthesis, but circFL-seq synthesized second-strand cDNA with an anchor primer instead of template switching of CIRI-long. In contrast, isoCirc produced rolling circles by RCA with circularized first-strand cDNA. The RCA strategy was able to generate longer reads with more rolling circles that benefit the detection of high-quality CS. However, as a trade-off, isoCirc produced fewer reads with the same sequencing depth, which raised sequencing costs and weakened its ability to detect and accurately quantify high-quality circRNAs. In addition, during the self-ligation of full-length synthesized first-strand cDNA of circRNA, truncated first-strand cDNA of circRNA and contaminated cDNA synthesized from residual linear RNAs may also be self-ligated, thus leading to false discoveries of circRNA.

Because of the high error rate of nanopore sequencing, the computational pipeline of all three methods calculated CS from tandem repeats of reads to refine low-quality sequences, followed by localization of CS. However, CS detection could be oversensitive for circRNAs themselves containing multiple internal repeats. For example, the famous miRNA sponge *CDR1as*, which showed a high expression level in HEK293 cells by RNA-seq, failed to be detected by isoCirc. In our *circfull* algorithm, the RG mode can directly align sequenced reads to the reference genome to construct full-length circRNA, which can avoid the misrecognition of CS. With *circfull*, *CDR1as* was identified to be the circRNA with the highest expression in HEK293 cells by circFL-seq. Additionally, CS may ignore reads that are less than two full circles, especially for RCRT-based methods that generate shorter length products than RCA. In contrast, RG mode was able to identify full-length circRNAs from reads with only one full circle. Thus, *circfull* detected 56% more full-length circRNA reads with fewer than two circles in our data.

Previous short-read RNA sequencing was used to quantify circRNAs based on BSJ-spanning reads, while circFL-seq with full-length reads was employed to finely quantify circRNAs at the isoform level. Thus, circFL-seq is able to identify differential internal AS events, helping to unravel the function of sequence variants. Another novel application of full-length circRNA sequencing is to identify f-circ, which has been found to play important roles in cancer pathogenesis^27^ and has become a promising biomarker for liquid biopsy^29^. Current methods^27,29–31^ validated the proposed f-circ by designing divergent primers on known gene fusion junctions and thus were inefficient and possibly ignored f-circ from unknown gene fusions. With cancer cell line data, circFL-seq showed its ability to high-throughput identify f-circ, which will promote our knowledge of f-circ.

Together, circFL-seq is able to identify and quantify full-length circRNAs and has the advantage of high read coverage and thus requires lower sequencing cost. Moreover, we succeeded in pooling multiple libraries in a large-scale sequencer, PromethION, which will further decrease circFL-seq cost to twice that of RNA-seq. Thus, our developed circFL-seq process is an effective and affordable high-throughput full-length circRNA sequencing method for screening functional circRNAs at the omics scale.

## Materials and Methods

### Cell culture and RNA isolation

The human cell lines HeLa, SKOV3, MCF7, VCaP, SH-SY5Y, HEK293T and HEK293 were used in this study. Cell lines were cultured in DMEM (Invitrogen) supplemented with 10% FBS (YEASEN) and 1% penicillin/streptomycin (Solarbio) at 37°C with 5% CO_2_. Cell lines were collected at 80 to 90% confluency. All cell lines were authenticated and tested negative for mycoplasma contamination. Total RNA was extracted by the FastPure Cell/Tissue Total RNA Isolation Kit (Vazyme) according to the manufacturer’s instructions. Total RNA from the human brain (#636530) and testis (#636533) was purchased from Clontech.

### circRNA enrichment for library construction of RNA-seq and circFL-seq

#### rRNA depletion

Similar to a previous method^32^, nonoverlapping synthetic DNA probes of complementary sequences (Supplementary Table 4) of 18S and 28S rRNA at a final concentration of 1 μM, as well as 5S and 5.8S rRNA, 12S and 16S mtrRNA, ETS, and ITS at a final concentration of 0.1 μM were pooled. One microliter of DNA probe was mixed with 2 μg of total RNA and 2 μL of hybridization buffer (750 mM Tris-HCl, 750 mM NaCl) at a final reaction volume of 15 μL. The mixture was heated to 95°C for 2 minutes, slowly cooled to 22°C (−0.1°C/s), and incubated for an additional 5 minutes at 22°C. Two microliters of thermostable RNase H (NEB) was added along with 2 μL of 10× RNase H buffer at a final reaction volume of 20 μL, incubated at 50°C for 30 minutes and placed on ice. DNA probes were removed by 2.5 μL DNase I (NEB) with 3 μL 10× DNase I buffer at a final volume of 30 μL, incubated at 37°C for 30 minutes and placed on ice.

#### Poly(A) tailing

The reaction above was then mixed with 4 μL ATP (10 mM), 4 μL 10× Poly(A) Polymerase Reaction Buffer and 1 μL Poly(A) Polymerase (NEB) for poly(A) tailing and incubated at 37°C for 10 min. Purified RNA was isolated by 2.5× RNA Clean Beads (Vazyme) according to the manufacturer’s instructions. *RNase R treatment*. For circFL-seq, RNA was incubated at 37°C for 30 min and 70°C for 10 min in a 10 μl reaction that contained 1 U RNase R (Lucigen) and 1 μl 10× RNase R buffer. For RNA-seq, RNA was incubated under the same conditions but in a 20 μl reaction containing 2 U RNase R and 2 μl 10× RNase R buffer.

### Full-length circRNA cDNA preparation for circFL-seq

#### Reverse transcription

After enrichment of circRNAs, collected RNAs were reverse transcribed into first cDNA strands in a 20 μl reaction by P1-N6 (5’-GTCGACGGCGCGCCGGATCCATANNNNNN-3’) with HiScript III reverse transcriptase (Vazyme) for 5 min at 25°C, 50 min at 50°C, 2 min at 70°C, and 5 s at 85°C, followed by purification with 0.75× DNA Clean Beads (Vazyme) according to the manufacturer’s instructions.

#### Poly(A) tailing

Then, poly(A) tails were added at the 3’ ends in a 20 μl reaction by terminal deoxynucleotidyl transferase (Invitrogen) with final dATP and ddATP concentrations of 2.5 mM and 25 μM, respectively, followed by purification with 0.75× DNA Clean Beads. *Second-strand synthesis*. Next, second-strand cDNAs were synthesized using P2-T24 (5’-ATATCTCGAGGGCGCGCCGGATCCTTTTTTTTTTTTTTTTTTTTTTTT-3’) by I-5 High-Fidelity DNA polymerase (MCLAB) at 98°C for 2 min, 50°C for 2 min, and 72°C for 5 min.

#### Amplification

Then, cDNAs were equally split into four 50 μl PCRs with primers P1 (5’-GTCGACGGCGCGCCGGATCCATA-3’) and P2 (5’-ATATCTCGAGGGCGCGCCGGATCC-3’) and amplified by 20 cycles of 98°C for 10 s, 67°C for 15 s, and 72°C for 75 s, followed by 0.5× DNA Clean Bead purification. Approximately 10–50 ng purified cDNAs were further amplified by 8–10 cycles in a set of four 50 μl PCR reactions with P1 and P2, followed by purification with 0.5× DNA Clean Beads.

### Nanopore library construction and sequencing

A DNA library for barcoding and ligation sequencing was prepared following protocols EXP-NBD104 and SQK-LSK109. Briefly, 0.5–1 μg of circFL-seq cDNA was repaired and dA-tailed by the NEBNext FFPE DNA Repair Mix (NEB) and NEBNext Ultra II End repair/dA-tailing Module (NEB), followed by purification with DNA Clean Beads. For multiplexing, repaired and end-prepped DNA was barcoded with Native Barcode by NEB Blunt/TA Ligase Master Mix (NEB), followed by purification with DNA Clean Beads. Barcoded samples were pooled together in equimolar amounts. Single sample without barcodes or pooled barcoded samples (700 ng) were ligated to an ONT Adapter with NEBNext Quick Ligation Module (NEB), followed by purification with DNA Clean Beads. The DNA library was mixed with sequencing buffer, and beads were loaded onto a PromethION or MinION R9.4 flow cell and run on a PromethION (performed by Grandomics) or MinION sequencer, respectively.

### cDNA library preparation and sequencing of RNA-seq

After circRNA enrichment, RNA was isolated by 2.5× RNA Clean Beads (Vazyme). cDNA libraries were constructed and barcoded by the VAHTS Universal V6 RNA-seq Library Prep Kit for Illumina (Vazyme) and VAHTS RNA Adapters set1/set2 for Illumina (Vazyme) according to the manufacturer’s instructions for the strand-specific rRNA depletion library. The cDNA library was fragmented, and an insert size of approximately 300 bp was selected. Eight libraries were pooled together and sequenced on the Illumina HiSeq X Ten platform of Annoroad Gene Technology with a paired-end read length of 150 bp.

### Computational analysis of full-length circRNAs with circFL-seq

Raw sequenced data in fast5 format files were transformed to fastq format files (kept reads with qscore ≥7.0) and were demultiplexed by guppy (4.2.2). We employed porechop (v0.2.4) to trim barcode and circFL-seq primers (P1, P2) and split chimeric reads to obtain clean reads for each sample. Then, RG mode of *circfull* (https://github.com/yangence/circfull) was employed to detect circRNAs with clean reads. The cRG mode of *circfull* was used to correct circFL-seq reads by *de novo* self-correction (DNSC) to calculate CS and the RG mode was rerun with a query sequence of three copies of CS. With primer sequences at both ends of sequenced reads, *circfull* detected strand origin of reads and adjusted circRNA results with strand information. Finally, *circfull* was used to filter out low-quality full-length circRNAs after integrating circRNA results.

#### RG mode

Clean reads were mapped to the human genome (hg19) by minimap2^33^ (v2.12) with the parameters ‘-ax splice –p 0.5’ or ‘-ax splice –p 0.5 -uf’ for transcript-strand reads. Aligned reads with chiastic overlapping segments were recognized as candidate circRNA reads (CCRs). CCRs were classified into three types: normal, fusion on the same chromosome, and fusion on different chromosomes. The boundaries of the chiastic segment of the CCRs were detected as potential BSJs. Forward splicing junctions (FSJs) were determined according to the skipped region from the reference. For each read, a pseudo-reference sequence was generated by concatenating two sequences from 150 nt upstream to 150 nt downstream of the BSJ region. Then, CCRs were realigned against the pseudo-references, and accurate sites of BSJs were determined by integrating multiple aligned BSJs, gene annotation (GENCODE v19) and canonical splicing motifs (GT/AG). FSJs of CCRs were also corrected based on the integration of multiple aligned FSJs, gene annotation and FSJs from other CCRs with the same BSJ. Next, the full-length circRNA was constructed based on the FSJ(s) and BSJ(s). Incorrect construction of full-length circRNAs caused by mistaken alignment against tandem repeat sequences of the reference genome was identified by TideHunter (v1.0) with parameters ‘-f 2 –c 1.2 −l’, and these circRNAs were filtered out.

#### The DNSC

CS of clean reads was detected by TideHunter (v1.0) with the parameters ‘-f 2 –c 1.5 -p 30 −l’. Following evaluation by Tandem Repeats Finder (v4.09) with parameters ‘2 5 7 80 5 5 2000 -h -ngs’, CS were removed if containing internal tandem repeats, defined by an alignment score >40 or length of internal tandem repeats longer than half of the CS.

#### cRG mode

To locate the genomic region of CS, pseudo-query sequences from three combined monomers of a CS were created. The position of full-length circRNA was detected with RG mode.

#### Identification of the strand origin of clean reads

CCRs with FSJs detected were first selected to determine the strand origin of sequenced reads, i.e., first or second strand, based on the GT/AG motif of the FSJ and mapped strand. For each 100 nt flanking end of the raw read, the maximum identical number of bases to primers P1, P2, T24 and their reverse complementary sequences was calculated by the Smith-Waterman algorithm. With these numbers of identical bases as predictor variables and strand origin as the target variable, a random forest classifier was trained to predict the strand origin of all clean reads. CircRNA with ambiguous strand direction was adjusted by rerunning RG mode with stranded reads.

#### Filtration of low quality circRNA

After RG detection supplemented with cRG detection, low-quality circRNAs with an unsplicing ratio of BSJ <0.1 were filtered out. CircRNA isoforms with reliable BSJs and FSJs were retained if more than half of the circRNA reads were perfectly aligned on ±4 bp of junctions. CircRNA BSJs were filtered out if both BSJs were located in ±30 bp of repeat elements.

### Reverse transcription for PCR and RT-qPCR validation

For PCR validation, sample RNA was reverse transcribed to cDNA products for 5 min at 25°C, 50 min at 50°C, 2 min at 70°C, and 5 s at 85°C with random hexamers by HiScript III. For RT-qPCR, sample RNA was reverse transcribed to cDNA products for 15 min at 37°C and 5 s at 85°C with random hexmers and oligo(T) according to the manufacturer’s instructions.

### Validation of full-length sequences of circRNA

A pair of divergent PCR primers was designed and included 6 to 8 additional G bases at the 5’ end (Supplementary file 5) to reduce the accumulation of short PCR products. With cDNA products of the reverse transcription or circFL-seq library as template, PCR amplification in a 25 μl reaction of Takara Ex Taq Hot Start was carried out by the following program: 98°C for 10 s followed by 3 cycles of 98°C for 10 s, 60°C for 30 s and 72°C for 90 s, then 30 cycles of 98°C for 10 s and 72°C for 90 s, with a final extension at 72°C for 60 s. The PCR products were analyzed on 1.2% agarose gels (TSINGKE) and the rolling circle bands were cut out and extracted, followed by TA cloning to a pEASY-T1 cloning vector (TransGen). Then, the clones with inserts were Sanger sequenced.

### Validation of AS in circRNAs

Total RNA of the HeLa cell line (1 μg) was treated with RNase R (4 U) in a 10 μl reaction and then reverse transcribed to the cDNA products. For each circRNA isoform from the same BSJ, divergent primers (Supplementary file 6) targeting but not across splicing junctions were used to validate the AS junction site in a PCR volume of 25 μl. The PCR products were analyzed on 1.2% agarose gels, and the target bands were cut out and extracted, followed by TA cloning and Sanger sequencing.

### Quantification of circRNA expression by RT-qPCR

Total RNA of HeLa and SKOV3 cell lines (1 μg) w/wo RNase R treatment (4 U) was reverse transcribed for RT-qPCR. For specific circRNAs, primers (Supplementary file 7) across a BSJ with cDNA as template in a 20 μl reaction were set according to the manufacturer’s instructions for ChamQ Universal SYBR qPCR Master Mix (Vazyme). For specific circRNA isoforms, we designed primers (Supplementary file 8) for both specific BSJs and alternative FSJs in the reaction. For the quantification of internal AS events, two major circular isoforms 1 and 2 from the same BSJ were selected for the evaluation of the transcript ratio with the following formula: 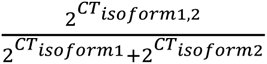. Primers targeting *GAPDH* as a reference gene were used. Thermal cycling was carried out on an Applied Biosystems 7500 Fast system at 95°C for 5 min, followed by 40 cycles of 95°C for 10 s and 60°C for 24 s.

### Validation of f-circ

Total RNA of the MCF7 cell line (1 μg) w/wo RNase R treatment (4 U) was reverse transcribed for PCR. We validated the full-length f-circ with primers (Supplementary file 9) for the two fusion junctions by RCRT and Sanger sequencing. Linear RNA was isolated to determine the origin of fusion junctions from BS or gene fusion. Because of the uncertainty of the poly(A) tail in linear RNA, total RNA was first treated with poly(A) tailing (NEB), and then linear RNA was selected by the poly(A) mRNA magnetic isolation module (NEB) according to the manufacturer’s instructions. PCR was performed with the same primers of both fusion junctions of f-circ. For quantification of the fusion junction of f-circ, total RNA of the MCF7 cell line (1 μg) w/wo RNase R treatment (4 U) was reverse transcribed for RT-qPCR. Primers (Supplementary file 10) of target fusion junctions were designed for RT-qPCR.

### CircRNA analysis from RNA-seq

RNA-seq data were aligned to the human reference genome (hg19) by BWA^34^ (v0.7.17-r1188). CircRNA BSJs were detected and quantified by CIRI2 with gene annotation (GENCODE v19). Full-length circRNA structures were constructed by CIRI-AS^35^, CIRI-full^10^ and CIRI-vis^36^. For fusion junctions, we directly searched the ± 10 bp of junction site in RNA-seq reads.

### Analysis of isoCirc data

isoCirc nanopore sequencing data were downloaded from the Sequence Read Archive (SRA: SRP235284). CircRNAs were analyzed with the isoCirc computational pipeline (https://github.com/Xinglab/isoCirc) with the human reference genome (hg19) and gene annotation (GENCODE v19).

### Identification of differentially expressed circRNA

CircRNA BSJs with at least 10 read counts in two or more RNA-seq samples and two or more circFL-seq samples of either the HeLa or SKOV3 cell line were kept to identify differentially expressed BSJs. DESeq2 with ‘mean’ fitType was employed to analyze differential expression between HeLa and SKOV3 cells. Differential BSJs at a FDR < 0.05 were recognized as DECs.

## Data availability

The circFL-seq and RNA-seq data produced by this study have been deposited in SRA (PRJNA722575). The information of circRNAs detected by circFL-seq is available in the figshare repository (https://doi.org/10.6084/m9.figshare.14265650.v1). The computational software *circfull* can be accessed from https://github.com/yangence/circfull.

## Acknowledgements

The work was supported by grants from Beijing Municipal Science and Technology Commission of China (7212065), Chinese Institute for Brain Research, Beijing (2020-NKX-XM-01), and Beijing Municipal Science and Technology Commission of China (Z181100001518005).

## Author contributions

Zelin Liu: Conceptualization; Resources; Data curation; Software; Formal analysis; Validation; Investigation; Visualization; Methodology; Writing - original draft Changyu Tao: Data curation; Investigation Shiwei Li: Resources; Validation Minghao Du: Resources; Data curation Yongtai Bai: Resources; Validation Xueyan Hu: Resources; Writing - review and editing Yu Li: Resources Jian Chen: Resources; Supervision Ence Yang: Conceptualization; Supervision; Funding acquisition; Writing - original draft; Project administration

## Competing interests

The authors declare no competing interests.

---

**Figure 1–figure supplement 1.**
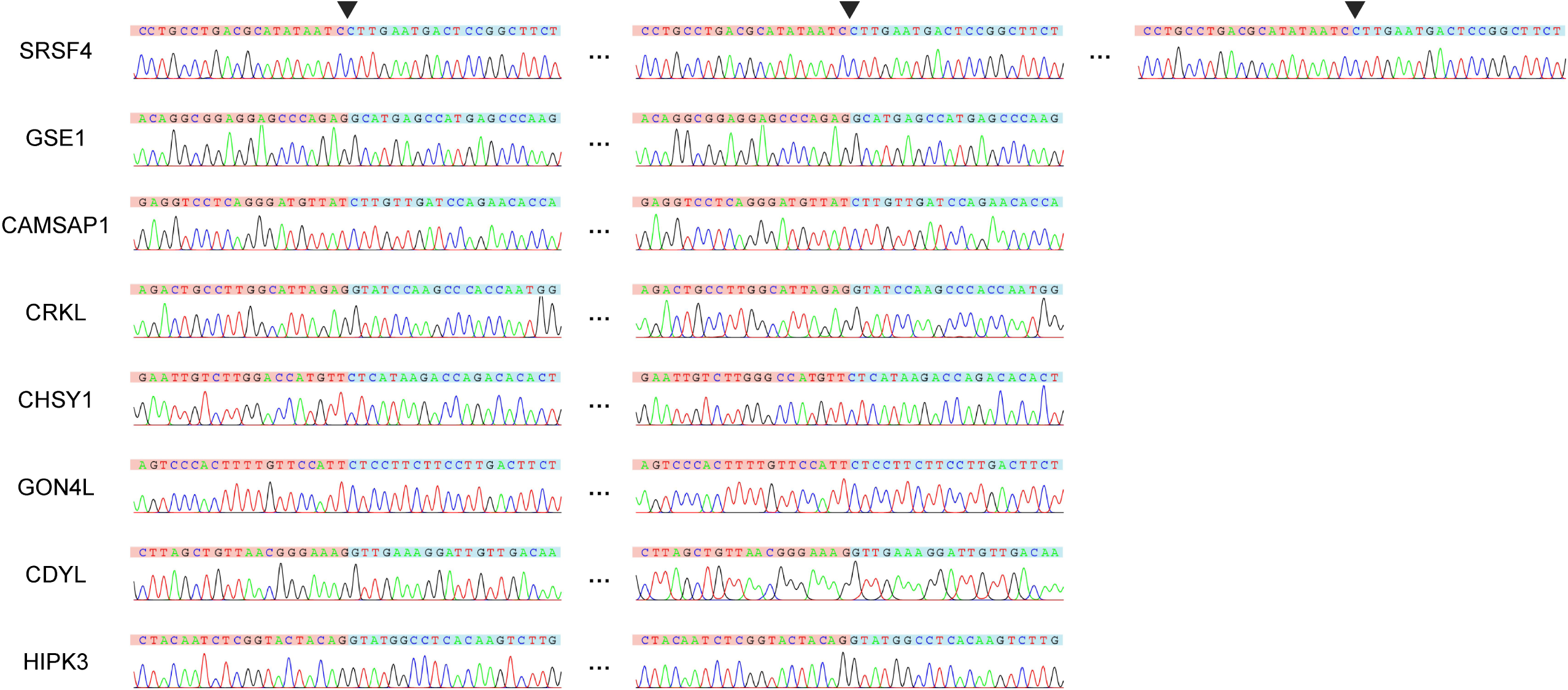
Sanger sequencing of rolling circular bands. PCR were performed with circFL-seq library of HEK293T cell as template.

**Figure 2–figure supplement 1.**
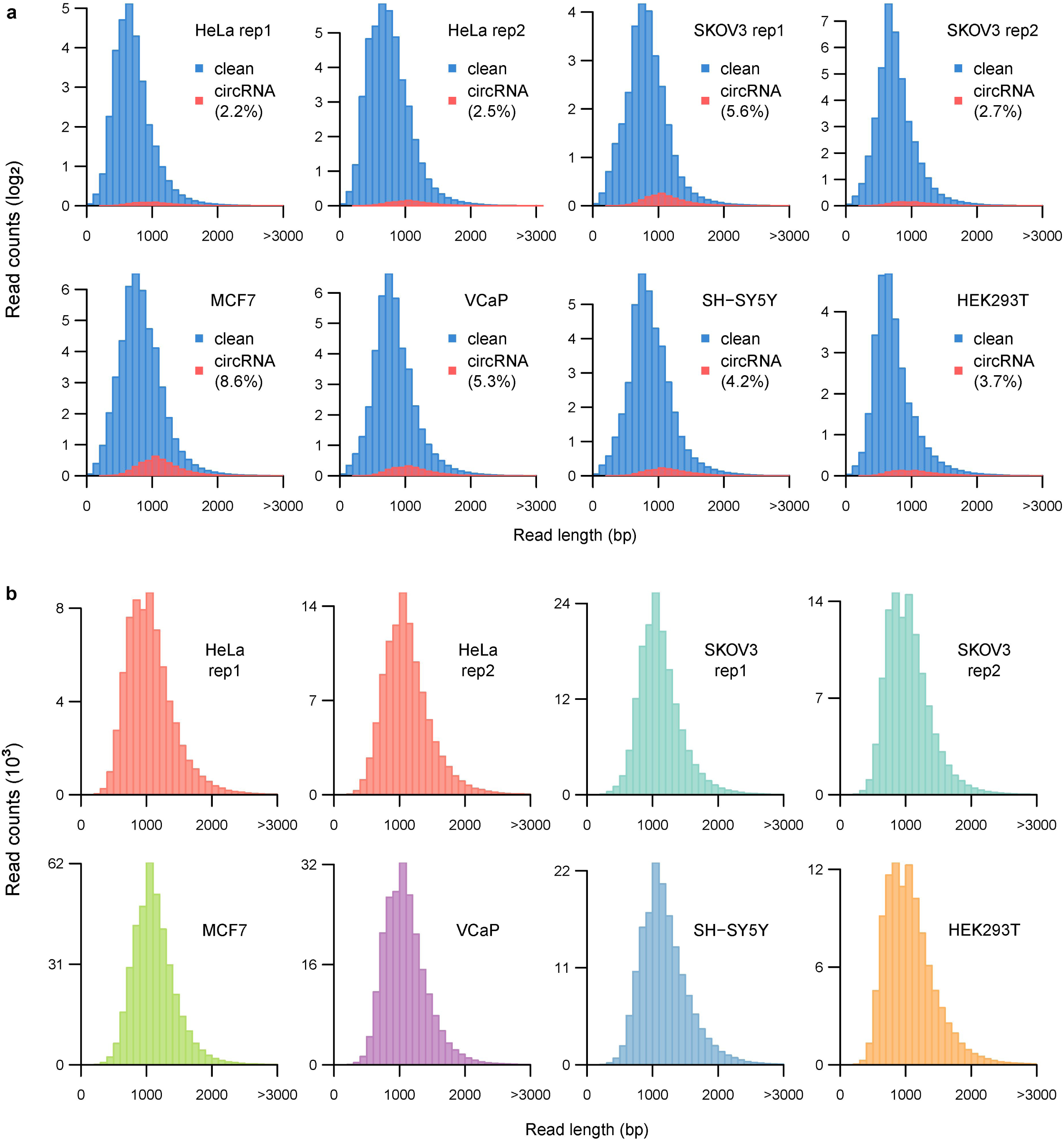
Clean reads distribution of circFL-seq data of six cell lines. **a.** Histograms showing the distribution of clean reads (blue) and full-length circRNA reads (red) for each sample. The percentages represent the circRNA reads proportion in clean reads. **b.** Histograms showing the distribution of full-length circRNA reads amounts.

**Figure 2–figure supplement 2.**
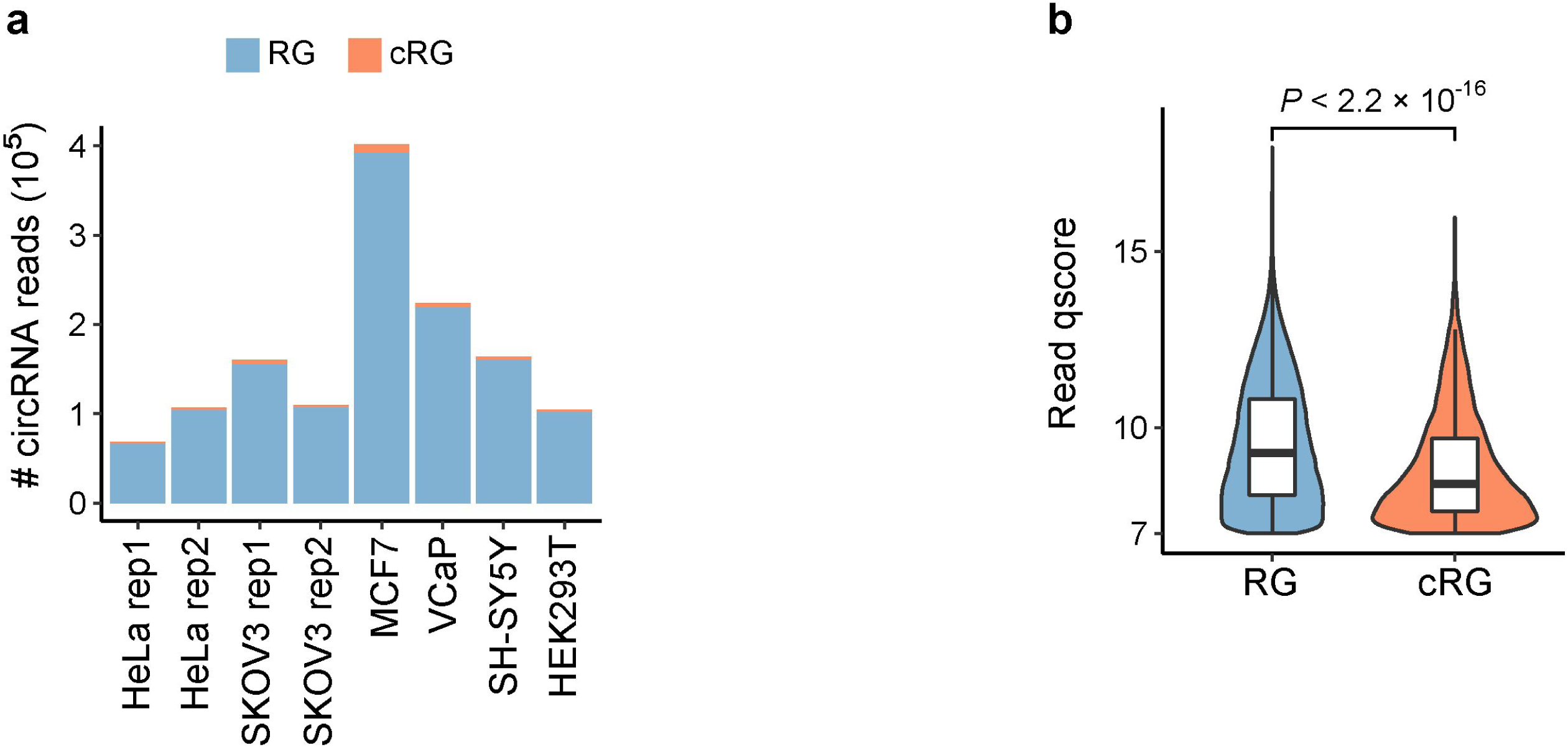
circRNA reads identified from circFL-seq data of six cell lines by *circfull*. **a.** Stacked bar plot representing the number of full-length circRNA isoforms detected by RG and cRG for eight samples. **b.** Boxplot showing read qscore distribution of circRNA isoforms of all samples. The qscore representing the read quality was extracted from the sequencing summary file.

**Figure 2–figure supplement 3.**
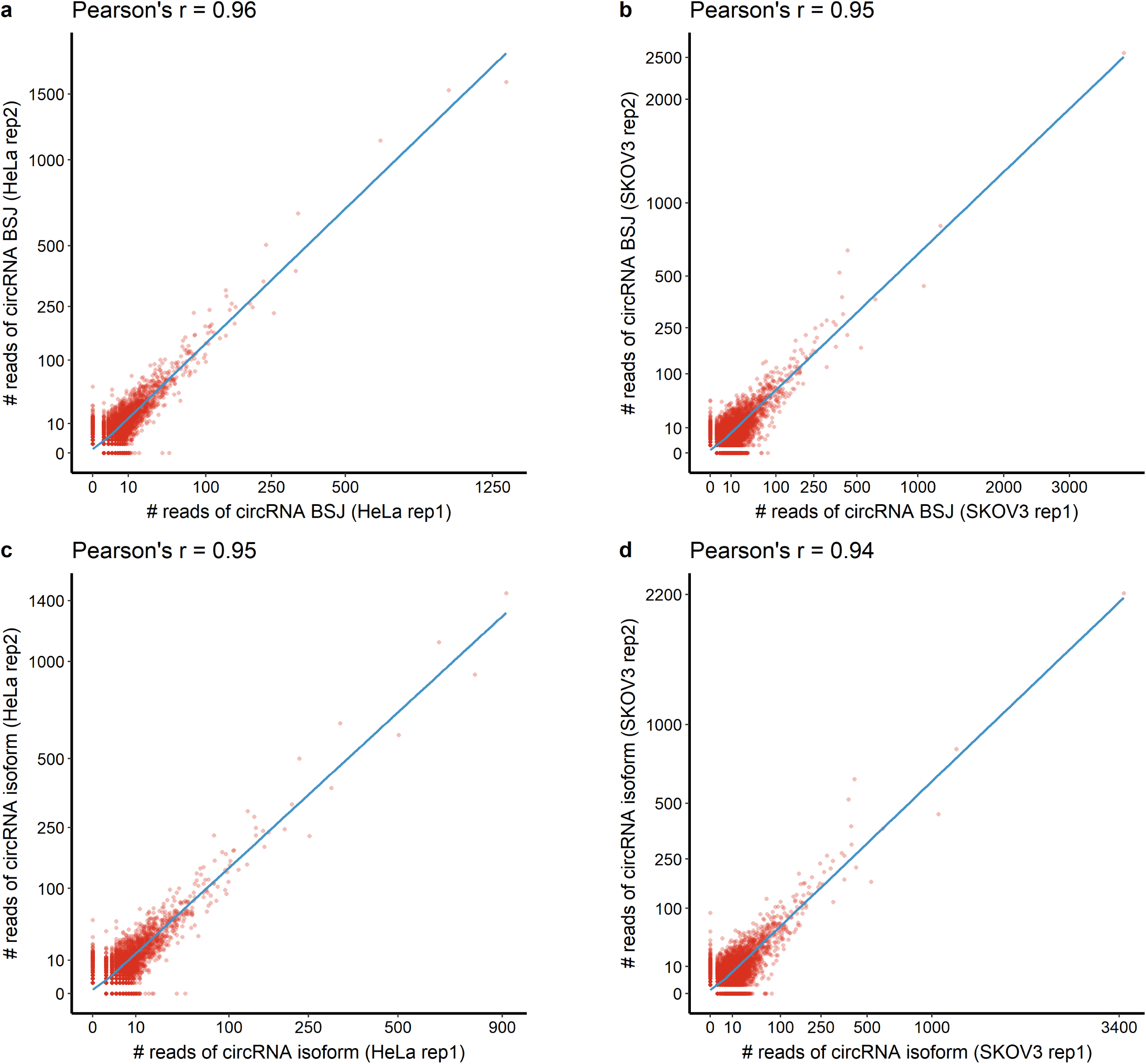
Scatter plot showing the correlation of circRNA at BSJ level (a,b) and isoform level (c,d) between circFL-seq replicates. CircRNA BSJs/isoforms with read counts >0 in at least one replicate were included.

**Figure 2–figure supplement 4.**
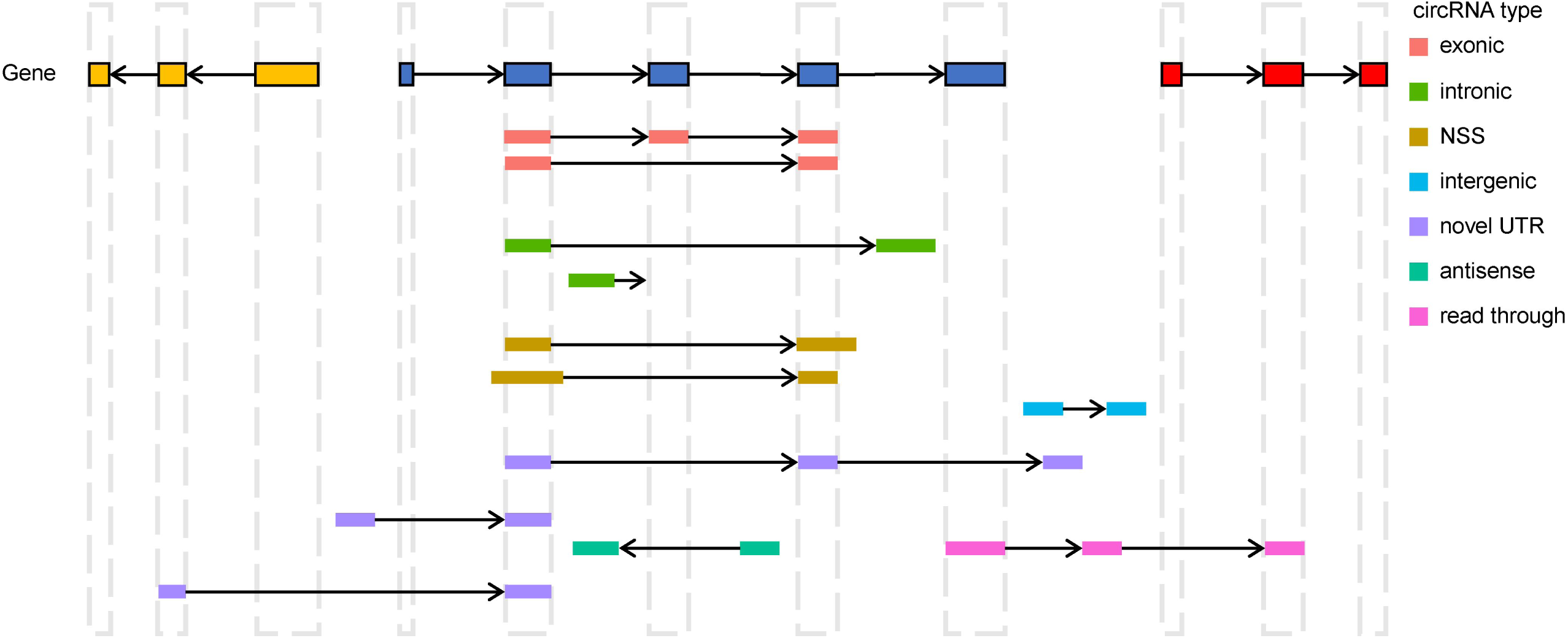
Diagram of circRNA types annotated by *circfull*. The classification is based on the positions of BSJs and boundary exons following below principles: **exonic:** CircRNA body is totally located inside of one gene from the same strand. Both of the BSJs are identical to annotated junctions. **intronic:** CircRNA body is also totally located inside of one gene from the same strand. But at least one of the boundary exons is not overlapping with any annotated exon. **novel splicing site (NSS):** CircRNA body is also totally located inside of one gene from the same strand. However, both boundary exons are overlapping with annotated exons, with at least one BSJ different to annotated linear junction. **intergenic:** The whole body of circRNA is located in intergenic region. **novel UTR:** CircRNA body partially overlaps with only one gene from the same strand, and at least one BSJ is located in intergenic region. **antisense:** There is no overlapping between circRNA and any gene from the same strand. But the circRNA overlaps gene(s) from the antisense strand. **read through:** BSJs are located in different genes with the same strand.

**Figure 2–figure supplement 5.**
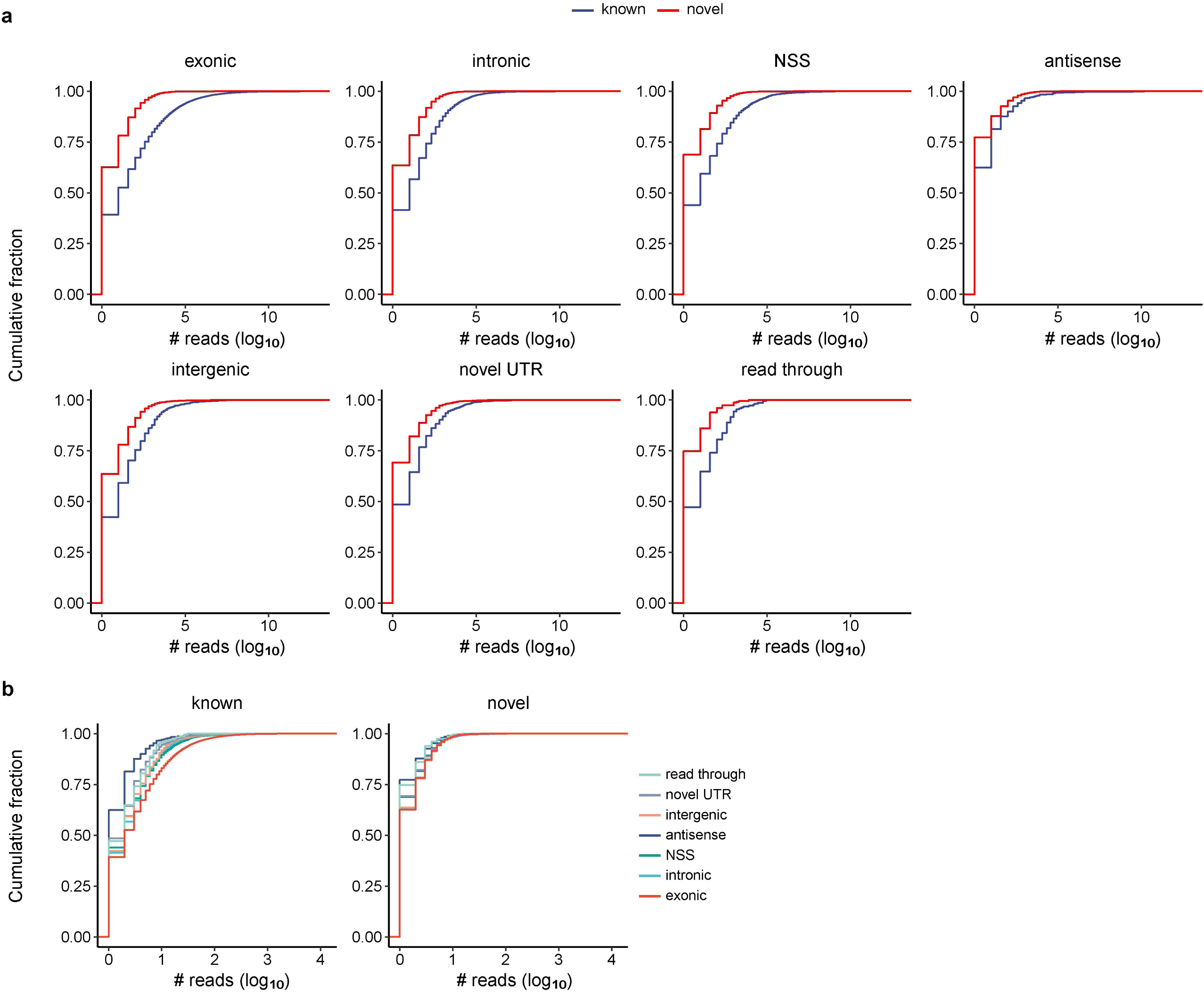
Cumulative distribution of read counts for circRNA isoforms identified by circFL-seq from six cell lines. CircRNA isoforms were classified based on database status and annotation types. **a.** For each annotation type (exonic, intronic, NSS, intergenic, novel UTR, antisense, read through), the cumulative distribution of read counts is classified to known and novel status based on their BSJs annotated in circRNA database or not. **b.** For known and novel status, the cumulative distribution of read counts is classified to seven annotation types.

**Figure 2–figure supplement 6.**
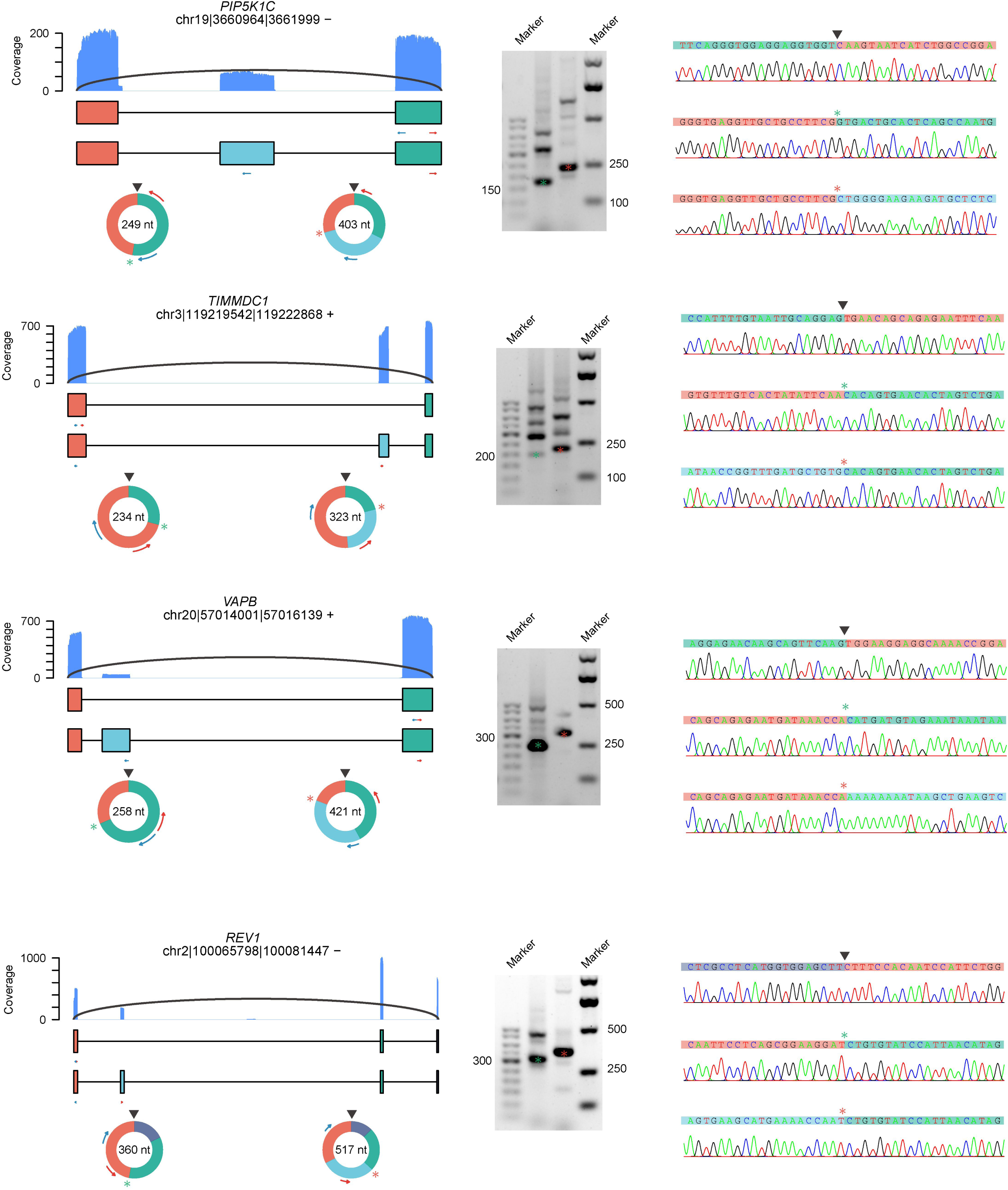
CircRNAs with exon skipping validated by RT-PCR and Sanger sequencing in HeLa cells. RT-PCR were performed with RNase R-treated RNA. Coverage of full-length circRNA reads mapped on reference genome were shown.

**Figure 2–figure supplement 7.**
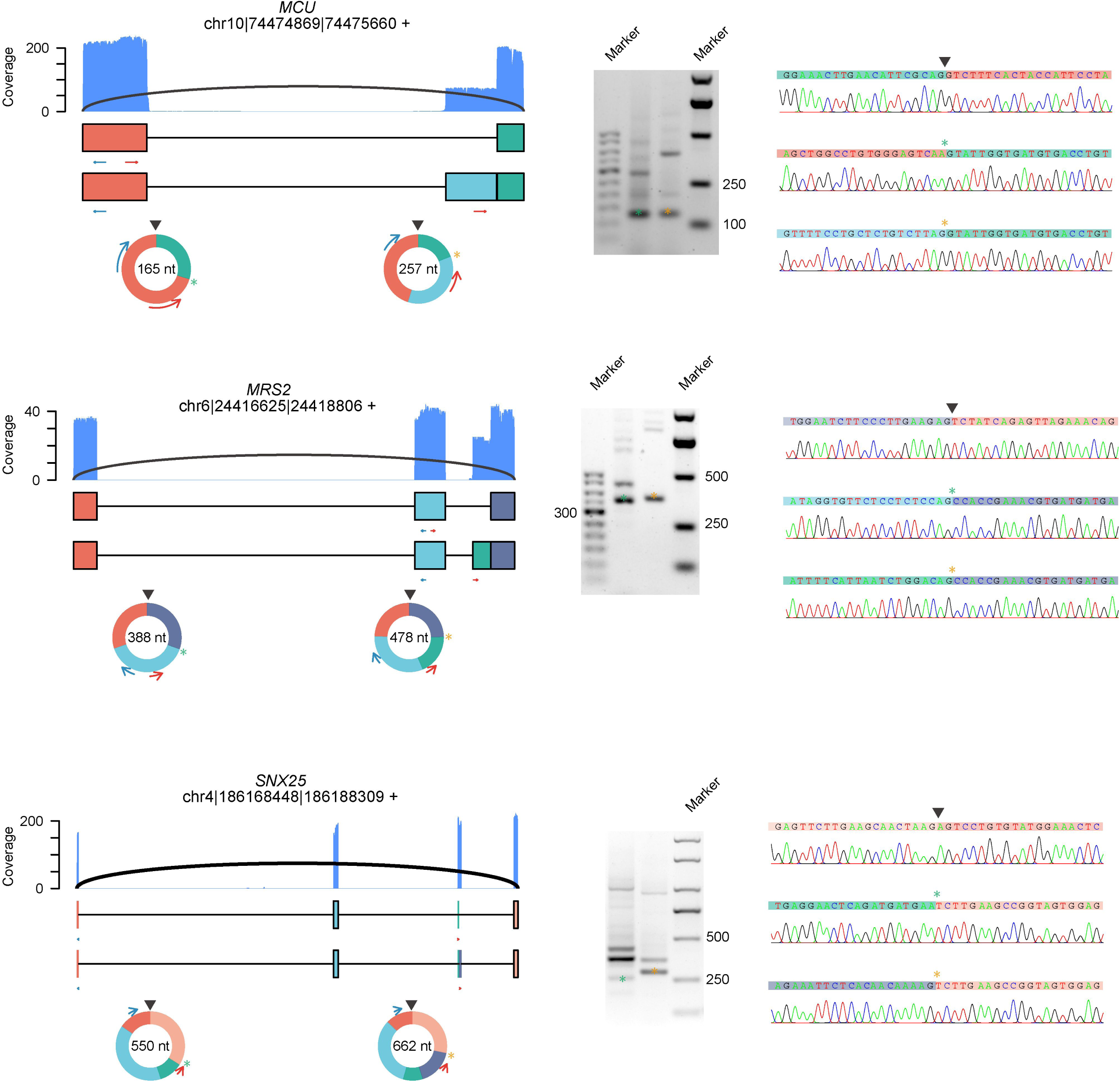
CircRNA with alternative 3’/5’ splicing site (A3SS for circRNAs from *MCU* and A5SS for circRNA from *MRS2*) validated by RT-PCR and Sanger sequencing in HeLa cells. RT-PCR were performed with RNase R-treated RNA. Coverage of full-length circRNA reads mapped on reference genome were shown.

**Figure 2–figure supplement 8.**
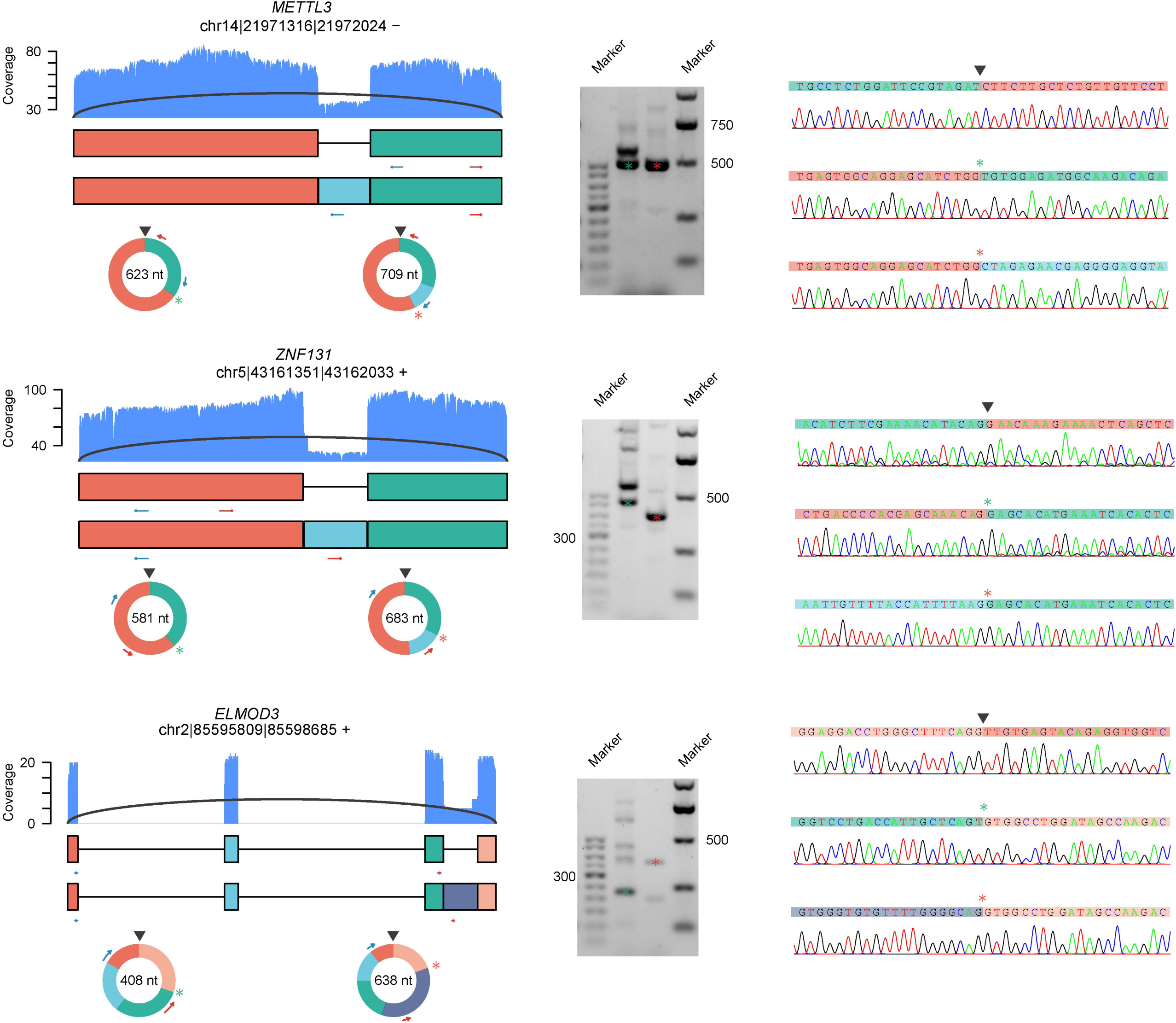
CircRNAs with intron retention validated by RT-PCR and Sanger sequencing in HeLa cells. RT-PCR were performed with RNase R-treated RNA. Coverage of full-length circRNA reads mapped on reference genome were shown.

**Figure 3–figure supplement 1.**
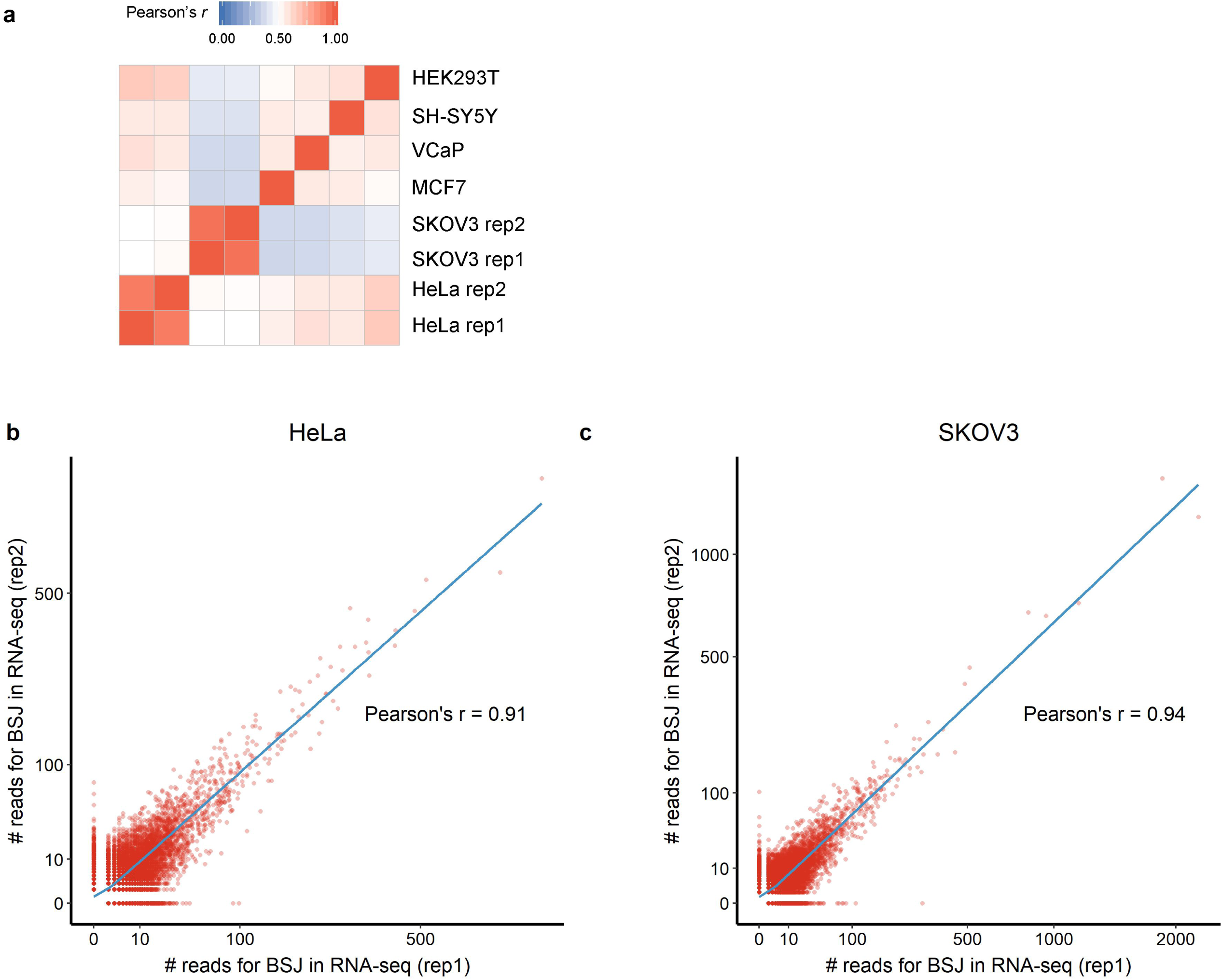
Correlations of circRNA BSJs among RNA-seq samples from six cell lines. **a.** Expression correlation matrix for circRNA BSJs among six cell lines. Color scale corresponds to Pearson’s correlation coefficients. Scatter plot showing the correlations of BSJs between HeLa (**b**) or SKOV3 (**c**) replicates. CircRNA BSJs with read counts >0 in at least one replicate were included.

**Figure 3–figure supplement 2.**
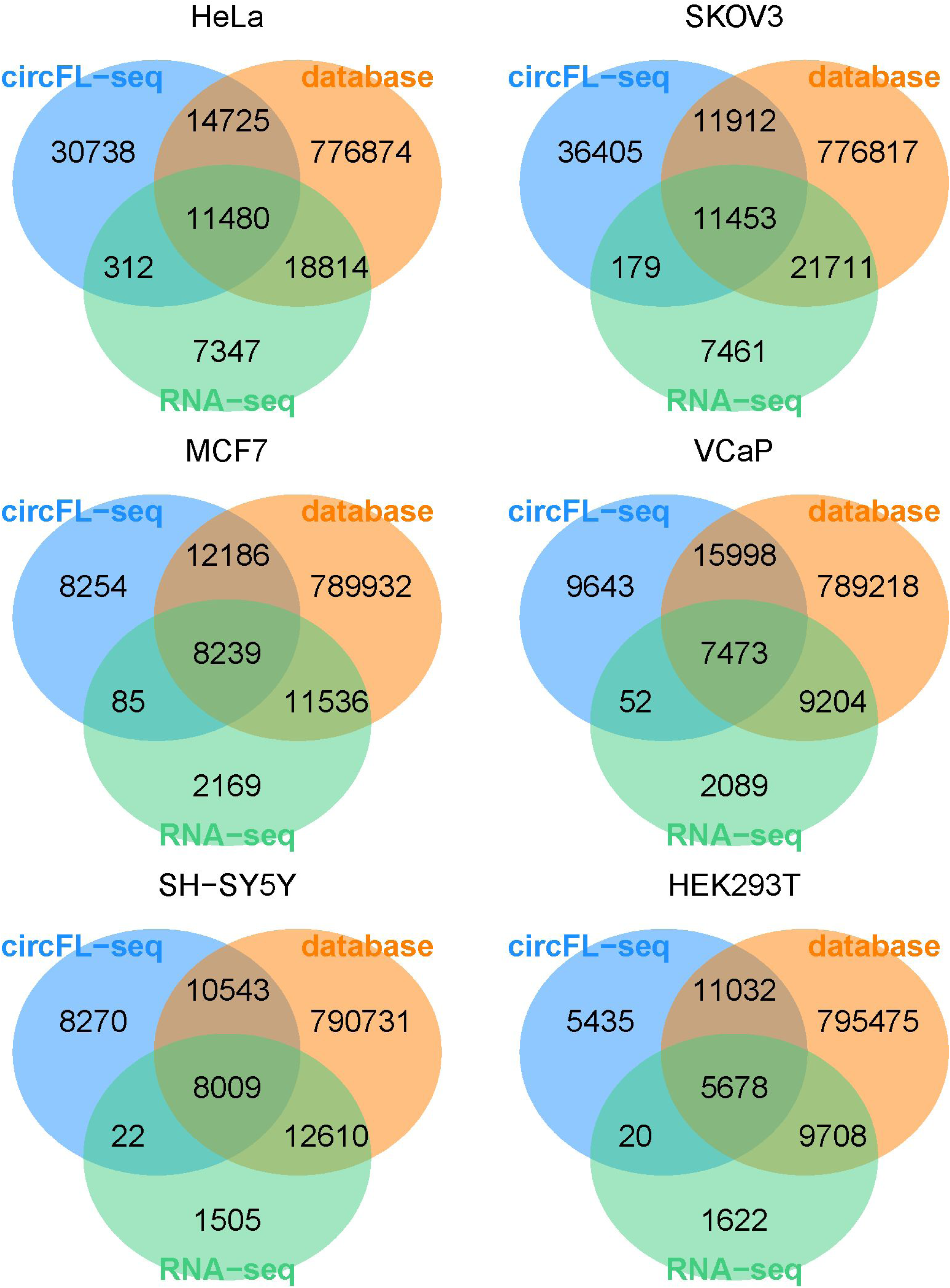
Venn diagram of BSJs detected by circFL-seq, RNA-seq, and database.

**Figure 3–figure supplement 3.**
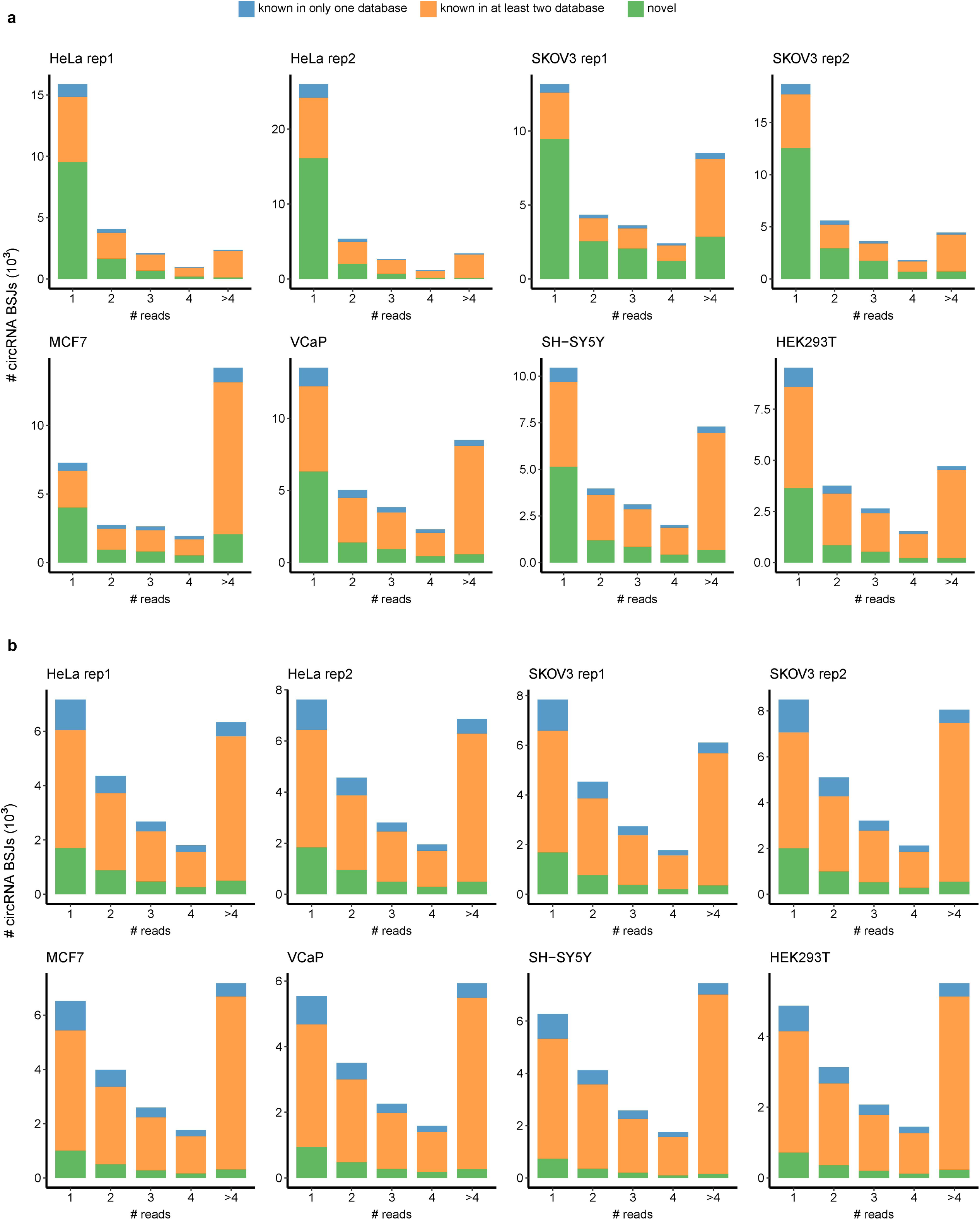
circRNA reads distribution of eight samples of six cell lines. Bar plot showing the distribution of known or novel circRNA BSJs with different read counts as threshold for circFL-seq (**a**) and RNA-seq (**b**) data.

**Figure 3–figure supplement 4.**
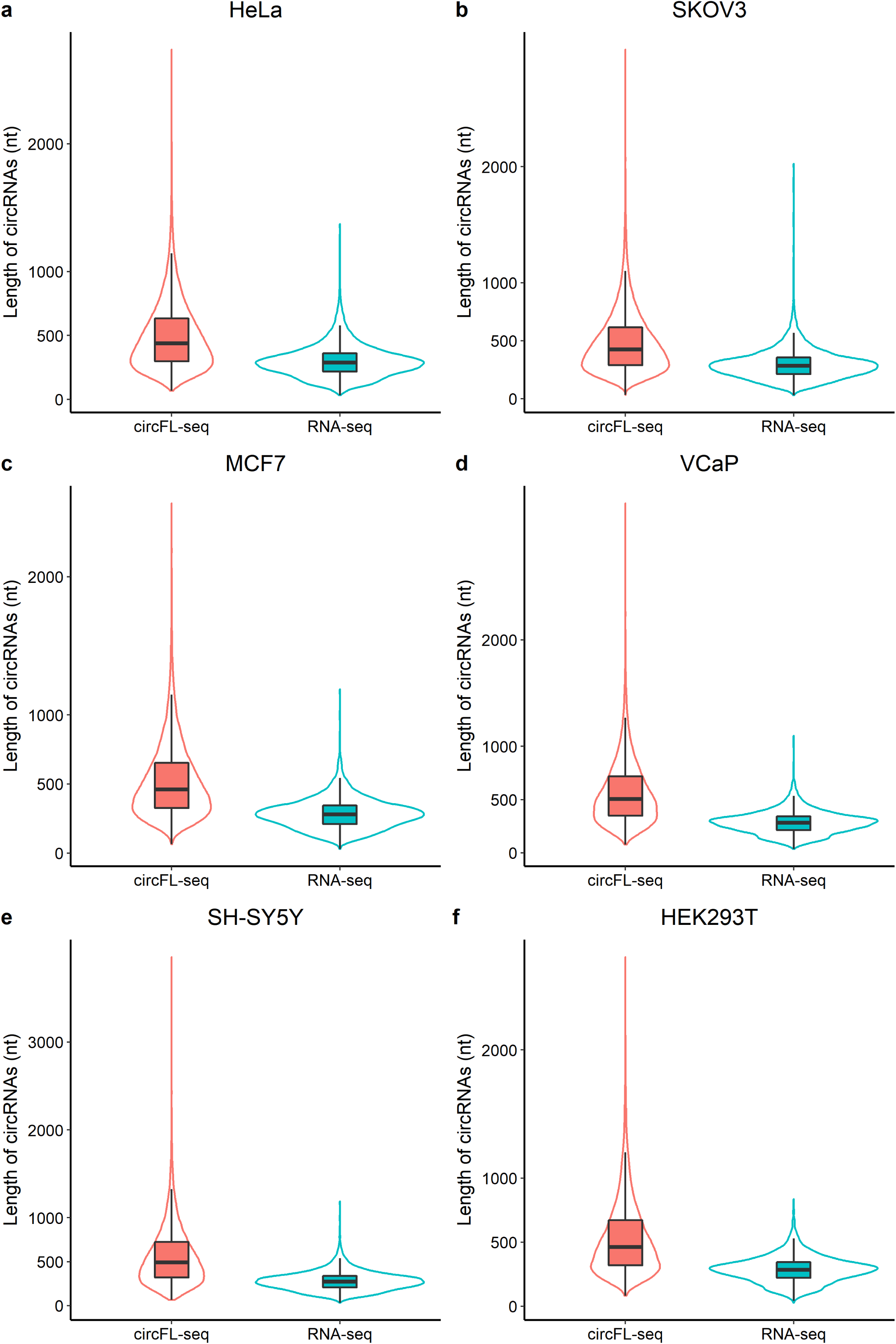
Comparison of circFL-seq and RNA-seq for length of full-length circRNA of six cell lines. For RNA-seq, full-length circRNAs were reconstructed by CIRI-full.

**Figure 3–figure supplement 5.**
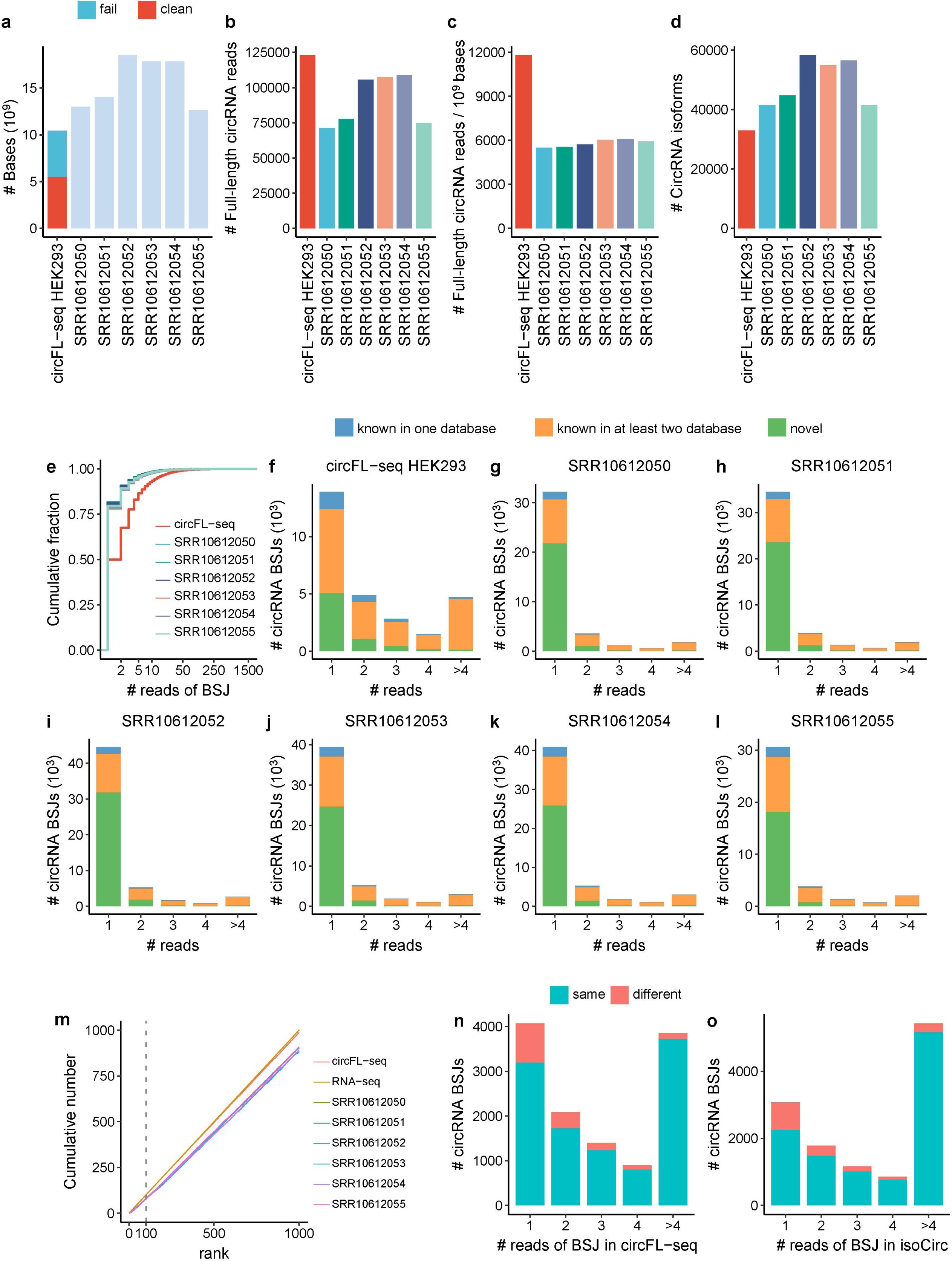
Comparison of circFL-seq and isoCirc for full-length circRNA detection in HEK293 cell line. **a.** Stacked bar plot showing the number of sequenced bases. circFL-seq includes fail and clean bases. Fail bases are from low quality reads with qscore <7 and trimmed adapters. **b.** Bar plot showing the number of full-length circRNA reads. **c.** Bar plot showing full-length circRNA reads per 10^9^ raw sequenced bases. **d.** Bar plot showing the number of full-length circRNA isoforms. **e.** Cumulative distribution of read counts of BSJs. **f-l.** Stacked bar plot showing the distribution of known or novel circRNA BSJs for different read counts. **m.** Plot showing the cumulative number for top expressed circRNAs of circFL-seq, isoCirc, and RNA-seq detected in database. Stacked bar plot showing the distribution of read counts of common circRNA BSJs detected in circFL-seq (**n**) and isoCirc (**o**). ‘same’ and ‘different’ represent BSJs w/wo same isoforms between circFL-seq and isoCirc. The isoCirc analysis in (**n,o**) combines circRNA results from all six HEK293 isoCirc libraries.

**Figure 3–figure supplement 6.**
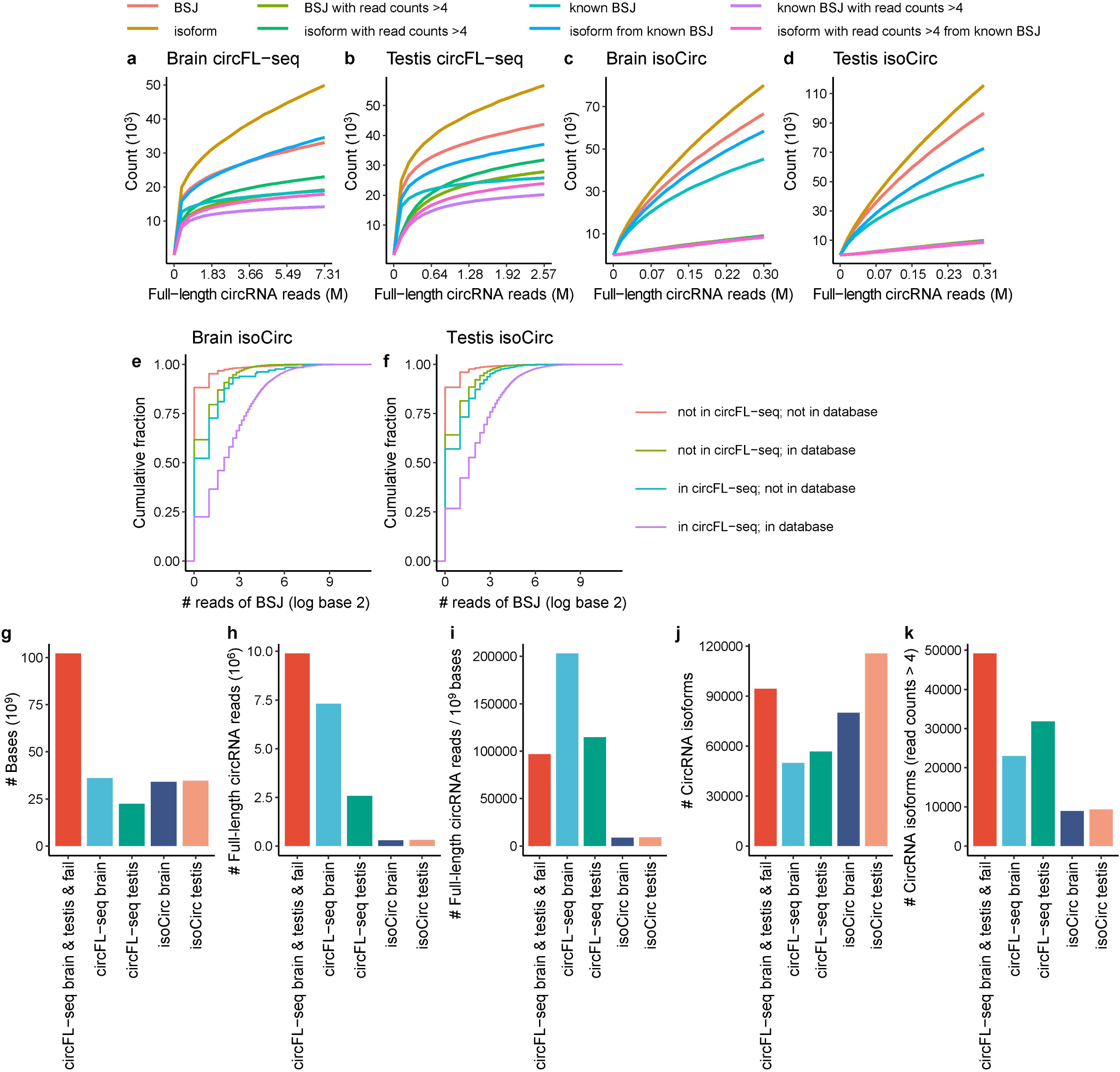
Comparison of circFL-seq and isoCirc for full-length circRNA detection in human brain and testis. **a-d.** Saturation curve of circRNA BSJs and isoforms for brain and testis data of circFL-seq (**a,b**) and isoCirc (**c,d**). Cumulative fraction of reads of BSJ of brain (**e**) and testis (**f**) data of isoCirc. **g.** Stacked bar plot showing the number of sequenced bases. Fail sequence data are from unclassified reads, low quality reads with qscore <7, and trimmed adapters. **h.** Bar plot showing the number of full-length circRNA reads. **i.** Bar plot showing full-length circRNA reads per 10^9^ sequenced bases. **j.** Bar plot showing the number of full-length circRNA isoforms. **k.** Bar plot showing the number of full-length circRNA isoforms with read counts >4.

**Figure 3–figure supplement 7.**
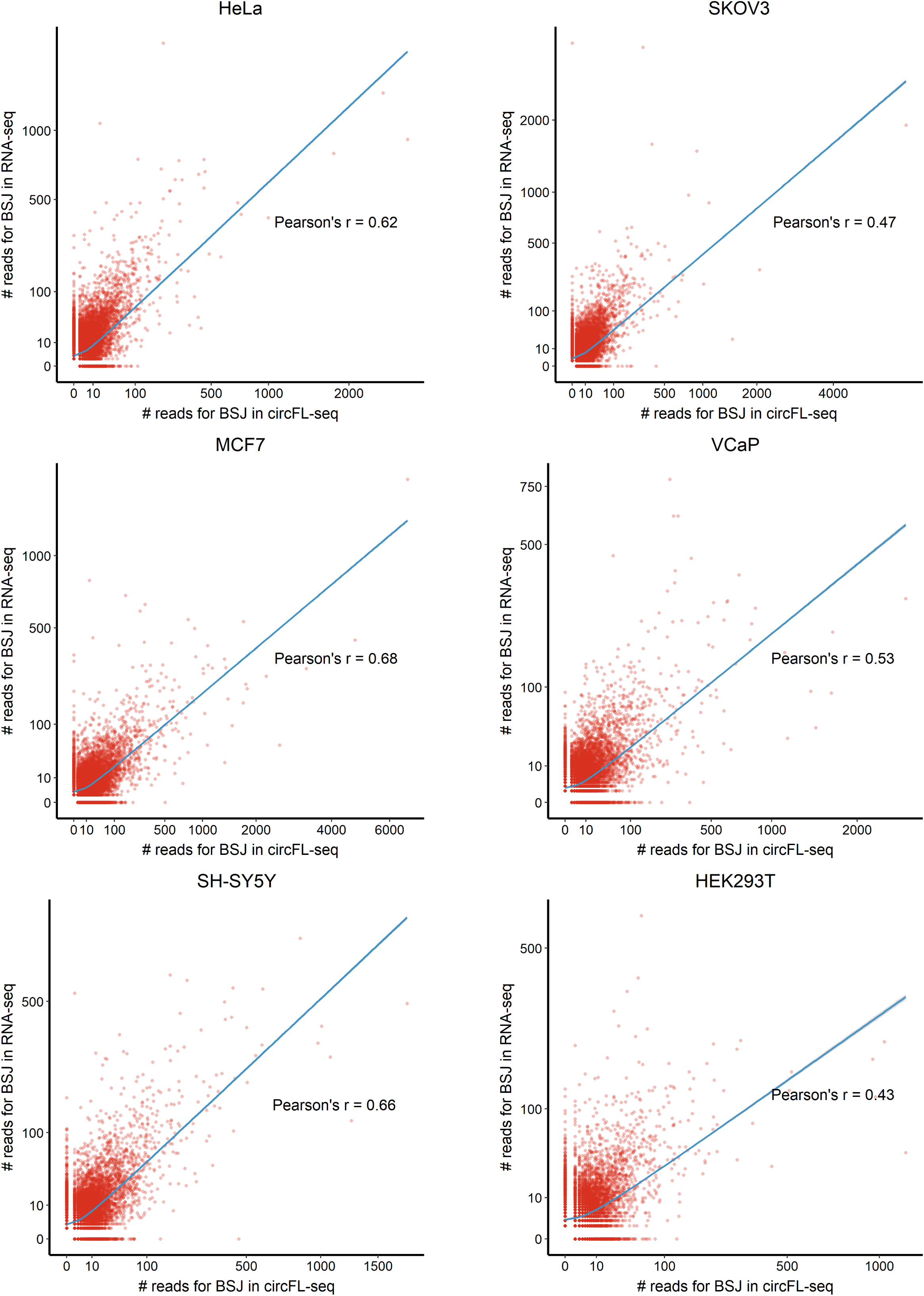
Scatter plot showing correlation of circRNA BSJs between circFL-seq and RNA-seq samples of six cell lines.

**Figure 3–figure supplement 8.**
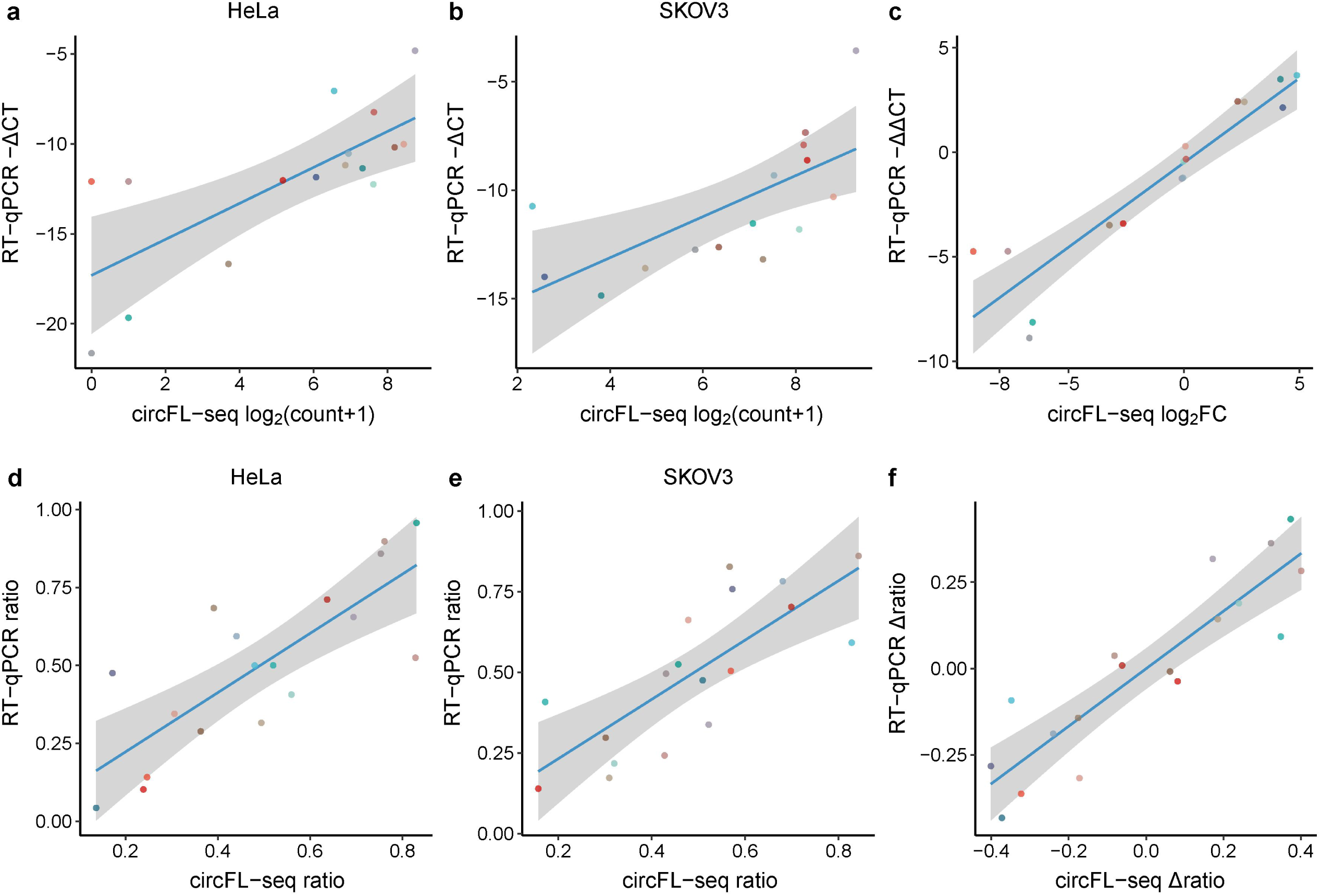
Evaluation of circRNA quantification between circFL-seq and RT-qPCR. The relative expression of target BSJs and isoforms quantified by RT-qPCR were performed with samples without RNase R treatment. GAPDH from total RNA without RNase R treatment was used as reference. **a,b.** Scatter plot showing the correlation of expression levels of 16 circRNA BSJs for HeLa (**a**) and SKOV3 (**b**) between circFL-seq and RT-qPCR. **c**. Scatter plot showing the correlation of fold change (log base 2) of the 16 BSJs for HeLa and SKOV3 between circFL-seq and RT-qPCR. **d,e.** Scatter plot showing the correlation of transcript ratio of 18 circRNA isoforms from 9 circRNA BSJs (each BSJ has two isoforms) for HeLa (**d**) and SKOV3 (**e**) between circFL-seq and RT-qPCR. **f.** Scatter plot showing the correlation of differential ratio (Δratio) of the 18 isoforms for HeLa and SKOV3 between circFL-seq and RT-qPCR.

**Figure 4–figure supplement 1.**
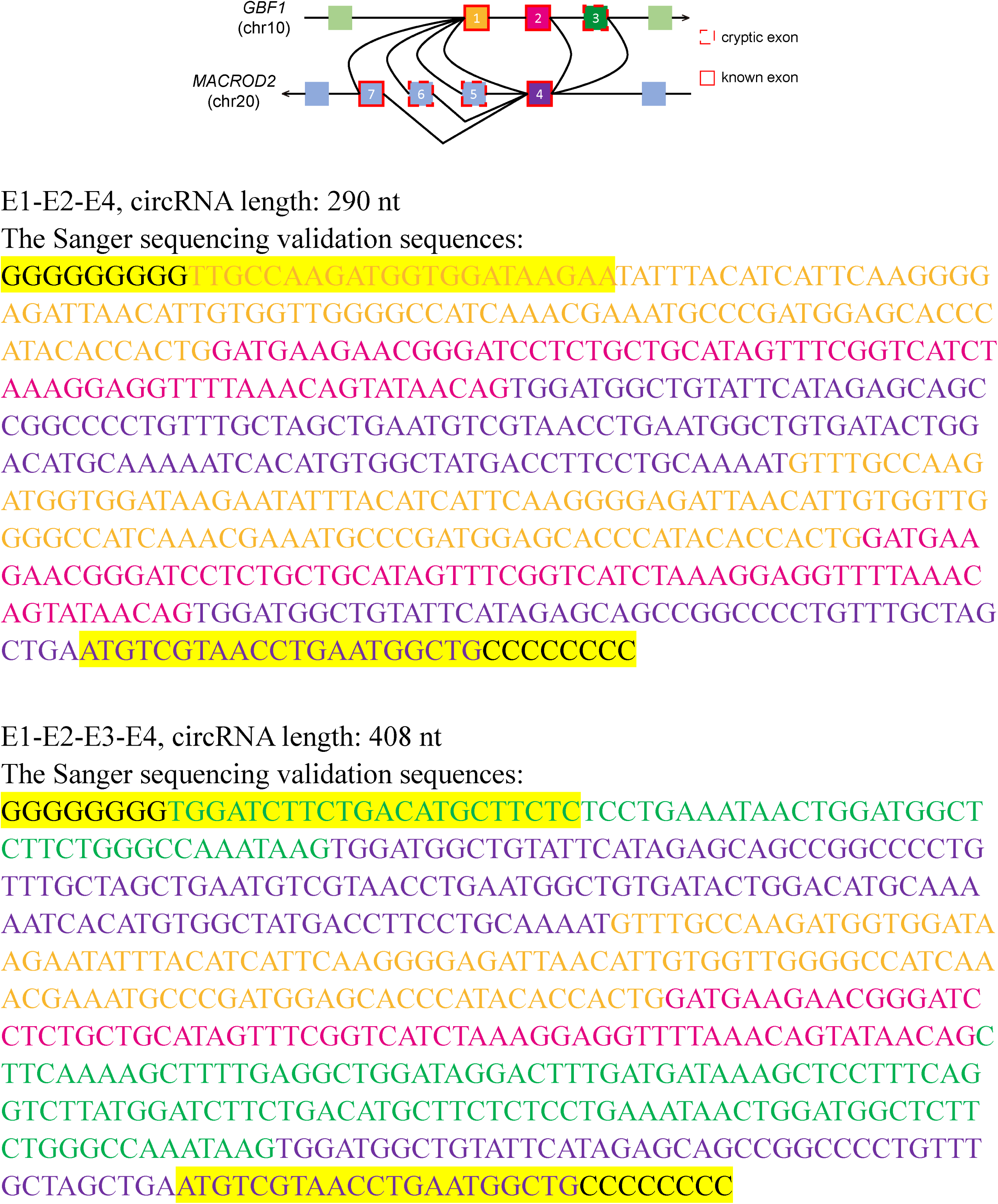
Sanger validation of sequences of f-circ from *GBF1:MACROD2.* The forward and reverse primers are highlighted.

**Figure 4–figure supplement 2.**
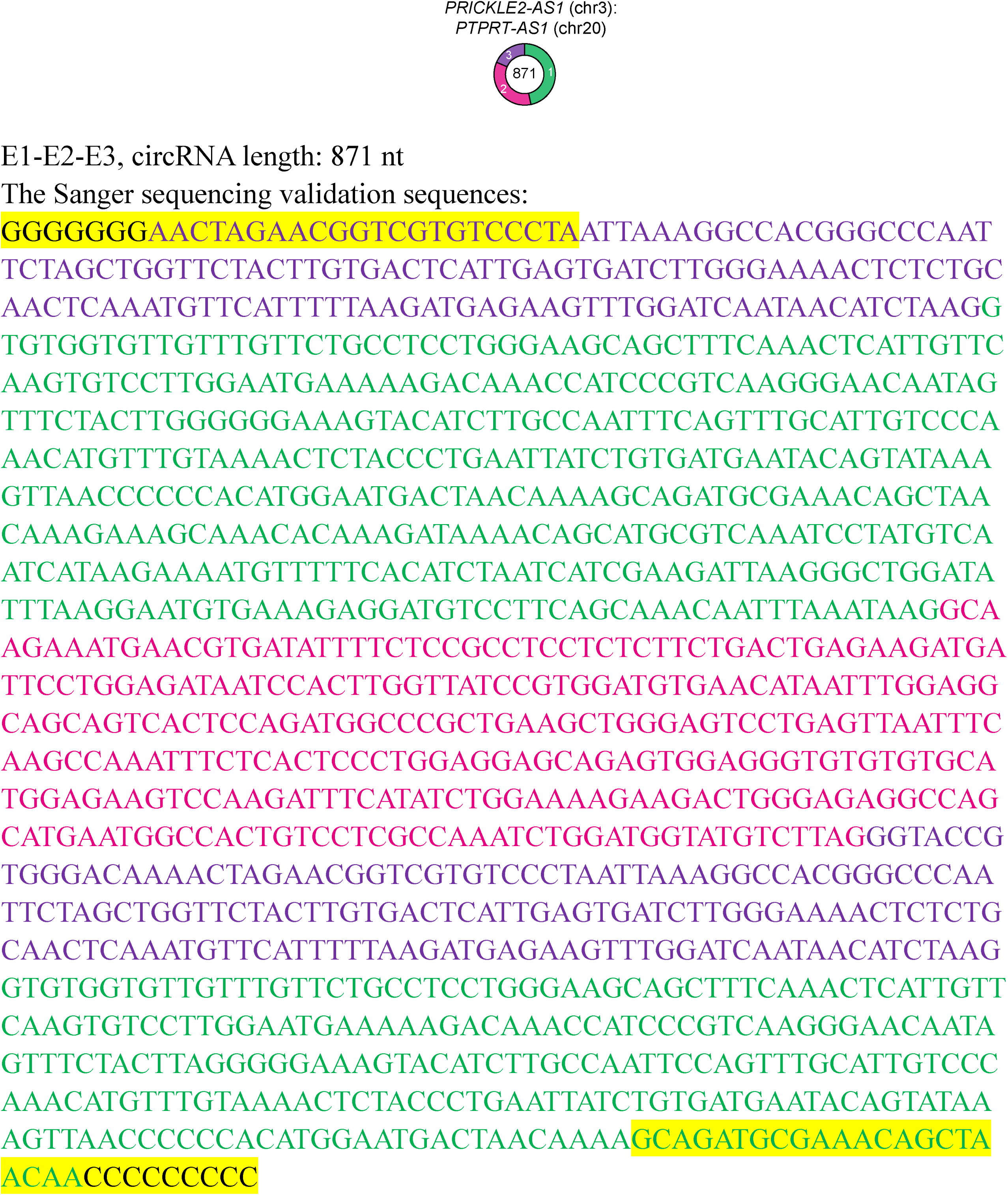
Sanger validation of sequence of f-circ from *PRICKLE2-AS1:PTPRT-AS1.* The forward and reverse primers are highlighted.

**Supplementary file 1. Data summary of circFL-seq library.**

**Supplementary file 2. Summary of alternative splicing events of circRNAs detected by circFL-seq.**

**Supplementary file 3. Data summary of RNA-seq library.**

**Supplementary file 4. Sequences of hybrid probes for rRNA degradation.**

**Supplementary file 5. Primers to validate rolling circles of circRNAs.**

**Supplementary file 6. Primers to validate alternative splicing of circRNAs.**

**Supplementary file 7. Primers to validate expression levels of circRNA BSJs by RT-qPCR.**

**Supplementary file 8. Primers to validate expression levels of circRNA isoforms by RT-qPCR.**

**Supplementary file 9. Primers to validate full-length sequence of f-circ.**

**Supplementary file 10. Primers to validate expression levels of f-circ junctions by RT-qPCR.**

## Notes

### Competing Interest Statement

The authors have declared no competing interest.

